# Prenatal diet buffers infant epigenetic changes linked to pollution and transient wheeze

**DOI:** 10.64898/2026.03.26.714555

**Authors:** Samantha Lee, Chaini Konwar, Robert Balshaw, Julia L. MacIsaac, Katia Ramadori, David T.S. Lin, Oscar Urtatiz, Kaja Z. LeWinn, Catherine J. Karr, Alicia K. Smith, Michael S. Kobor, Kecia N. Carroll, Nicole R. Bush, Meaghan J. Jones

**Author notes:** **Corresponding author:** Dr. Meaghan Jones, **Email:**. **Author Contributions:** SL conducted statistical analyses and wrote the first draft of the manuscript, CK performed data quality control, preprocessing, and estimation of cell type and genetic ancestry principal components, JLM, KR, DTSL, and OU performed nucleic acid extraction and microarray analysis of cord blood samples, RB provided guidance in statistical analysis. CJK estimated air pollutant exposures. NRB, KZL, KNC, and AKS supervised and coordinated sample and data collection and management in the CANDLE study. AKS, AJK, MSK, KNC and NRB provided guidance on study design and data analysis and reviewed manuscript. MJJ conceptualized the study, supervised data analysis, investigation, and reviewed and edited manuscript. All authors reviewed the final manuscript. **Competing Interest Statement:** The authors have no competing interests to declare.

## Abstract

Prenatal air pollution exposure is associated with childhood asthma, particularly among biological males. The mechanisms remain unclear, but may involve lasting epigenetic changes, such DNA methylation (DNAm), that occur during gestation in response to oxidative stress and inflammation. Higher maternal intake of “protective” micronutrients, like antioxidants, could buffer pollution-induced oxidative stress and inflammation to mitigate potentially adverse DNAm differences contributing to asthma. Using data from 515 CANDLE participants, we examined associations between prenatal NO_2_, PM_2.5_, and PM_10_ and cord blood DNAm, evaluated DNAm mediation of pollution associations with childhood wheeze phenotypes (transient, persistent, and late-onset), and assessed buffering of DNAm by maternal polyunsaturated fatty acid, vitamin C, or folate intake, and overall diet quality measured by the Alternative Healthy Eating Index-Pregnancy (AHEI-P). We identified 19, seven, and five regional DNAm differences associated NO_2_, PM_2.5_, and PM_10_. Mediation analyses suggested a role for *HLA-DPA1/DPB1* DNAm in NO_2_ and PM_2.5_ associations with transient wheeze. To assess buffering, we fit pollutant-by-diet interaction models, defining buffering as an interaction opposite in sign to the main pollutant effect. One or more micronutrients or AHEI-P attenuated pollutant effects at 16 of 19 NO_2_-associated DMRs, including *HLA-DPA1/DPB1*, and all PM_2.5_- and PM_10_-associated DMRs. However, attenuation of *HLA-DPA1/DPB1* DNAm did not significantly reduce the indirect effect of NO₂ on transient wheeze. In sex-stratified analyses, biological males exhibited lower PM_2.5_-associated DNAm in *SERPINB9*, a gene linked to lung function. These findings suggest prenatal air pollution alters DNAm, which may contribute to transient wheeze, with some differences partially buffered by maternal diet.

**Significance Statement:** Prenatal air pollution exposure contributes to child wheeze and asthma, potentially through the oxidative stress response and subsequent changes to infant DNA methylomes. Here, we used data from the CANDLE cohort to identify cord blood DNAm differences associated with NO_2_, PM_2.5_, or PM_10_. We examined if any alterations mediated the relationship between prenatal air pollution exposures and transient, persistent, or late-onset wheeze at age 4 to 6 years. Some of these DNAm differences appeared to be at least partially buffered by maternal micronutrients and/or overall diet quality.

## Introduction

Ambient air pollution is a heterogeneous mixture of gaseous and particulate matter (PM) compounds and is a widespread environmental health hazard. PM is further classified according by aerodynamic diameter, with particles <2.5µm (PM_2.5_) and <10µm (PM_10_) receiving the most attention as they can penetrate deep into the lungs and enter the periphery(1, 2). Here, transition metals on the surface of PM can catalyze oxidative reactions(3) and activate antioxidant and inflammatory pathways(4, 5), resulting in oxidative stress and inflammation. Nitrogen dioxide (NO_2_) is frequently used as a surrogate measure for overall air pollution exposure in epidemiological studies, but it also directly contributes to oxidative stress and inflammation, especially within the lungs(6, 7), where it oxidizes proteins and lipids, depletes antioxidants, activates cellular pathways involved in cell differentiation and death, and increases the release of inflammatory cytokines(7, 8). These effects can increase susceptibility to, or exacerbate, inflammatory conditions such as asthma(9, 10). Numerous investigations have reported positive associations between prenatal exposures to PM and NO_2_ air pollution and childhood asthma, with biological males appearing to be at especially high risk(11–13).

Childhood asthma is a complex condition in which wheezing is a common phenotype. Wheeze in children can be classified as no/infrequent wheeze, transient wheeze, persistent wheeze, and late onset wheeze(14). A systematic review identified a limited number of studies investigating the relationship between prenatal air pollution exposures and specific wheeze phenotypes, with some reporting associations with transient, persistent, and late-onset wheeze; however, findings varied across studies, likely due to differences in statistical methodology, wheeze and asthma definitions, and/or genetics(15). Additionally, these analyses did not include sex-stratified investigations, so it remains unclear whether susceptibility to these phenotypes differs by sex following prenatal air pollution exposure. While the exact molecular mechanisms connecting prenatal air pollution exposure to child wheeze and asthma remain unclear, growing evidence suggests it may operate through the biological embedding of *in utero* and early life oxidative stress and inflammation.

Biological embedding refers to a form of cellular “memory” of past experiences or exposures that can influence later biological responses and ultimately, health outcomes(16). This process is hypothesized to involve epigenetic modifications, including DNA methylation (DNAm) (16, 17), which refers to the addition of a methyl group to 5’ position of a cytosine in a cytosine-guanine dinucleotide (CpG). Previous investigations have identified relationships between prenatal air pollution exposures, altered DNAm at birth, and child health outcomes. For example, one study linked prenatal PM_2.5_ exposure to altered *UBE2S* DNAm at birth, with this DNAm difference positively correlated to asthma at six years of age(18). A meta-analysis reported associations between DNAm of the asthma-associated genes *NOTCH4* and *FAM13A* in cord blood, prenatal PM_10_ exposure, and gene expression in peripheral blood during adolescence(19). More recently, we identified *TRIM41* DNAm differences associated with prenatal NO₂ exposure that may contribute to atopy in male children within the CHILD study(20). Collectively, these studies have increased our understanding of the molecular pathways connecting prenatal air pollution exposure to child health outcomes, while highlighting key opportunities for more comprehensive investigation. Specifically, previous studies have not examined how air pollution-associated DNAm differences relate to distinct wheeze phenotypes, leaving important heterogeneity in respiratory outcomes unexplored. Additionally, most participants in these studies were mainly of European ancestry and from socioeconomically advantaged populations, potentially limiting the generalizability of findings to more diverse populations who may have higher asthma-associated prevalence and morbidity(21) and underscoring the need to determine which DNAm alterations are robust across human populations. Finally, no studies have explored potential buffers to reduce or prevent the effects of prenatal air pollutants on cord blood DNAm, representing an opportunity to help inform interventions aimed at lowering childhood asthma risk.

Given that oxidative stress and resulting DNAm differences appear to play an important role in mediating the health effects of prenatal air pollution exposure(22, 23), reducing oxidative stress could help prevent or attenuate DNAm alterations, which could weaken the association between prenatal air pollution exposure and childhood wheeze and asthma by reducing DNAm-driven disruptions in cellular function(24). Dietary antioxidants such as vitamin C can counteract oxidative stress, but are themselves depleted after prolonged exposure to oxidizing agents. For example, cigarette smoke (a type of air pollutant) is associated with lower plasma vitamin C levels in both smokers and infants born to mothers who smoked during pregnancy(24). A single-blind crossover intervention trial demonstrated that B-vitamin (folic acid, B6, B12) supplementation attenuates increases in inflammatory white blood cells after PM_2.5_ exposure in adults(25). In the CANDLE (Conditions Affecting Neurocognitive Development and Learning in Early Childhood) cohort, higher maternal plasma folate attenuated the negative association between prenatal PM_10_ exposure and child IQ measured at ages four to six (26). In another study, maternal vitamin C supplementation mitigated cord blood DNAm differences associated with prenatal cigarette smoking(27), with improved child respiratory outcomes, with higher lung function and decreased wheeze at age five years compared to the placebo group(28). Beyond individual nutrients, overall maternal diet quality may also influence oxidative stress and related biological pathways. One approach to measuring maternal diet quality is through the Alternate Healthy Eating Index for Pregnancy (AHEI-P), which captures intake of foods and micronutrients associated with lower chronic disease risk and those relevant to maternal and fetal health, including vitamin C, polyunsaturated fats, and folate(29). Despite this evidence, no study to date has delineated how overall maternal diet quality or micronutrient intake may modify the relationship between maternal prenatal air pollution exposure, infant DNAm at birth, and child respiratory outcomes including wheeze and asthma.

The present study examines whether maternal intake of protective micronutrients or overall diet quality can buffer the effects of prenatal air pollution exposure on cord blood DNA methylation (DNAm) and, in turn, influence the relationship between air pollution and childhood wheeze phenotypes at four years of age. Using the socioeconomically and racially diverse CANDLE cohort, we first conducted our main epigenome-wide analyses to identify individual CpG and regional DNAm differences associated with prenatal exposure to NO_2_, PM_2.5_, and PM_10_. We conducted secondary analyses, repeating the main analyses with biological sex interaction terms to explore whether DNAm differences varies between males and females to provide greater insight into the increased susceptibility of male children to wheeze and asthma with prenatal air pollution exposure. We also conducted secondary analyses, including air interaction terms for maternal intake of vitamin C, folate, or polyunsaturated fatty acids or overall maternal diet quality measured by AHEI-P score, to examine buffer buffering of DNAm differences identified in our main analysis. Finally, for DNAm differences that were associated with wheeze phenotypes, we explored the significance of the buffering effect of maternal micronutrient intake and overall diet quality using moderated mediation models.

## Materials and Methods

### Study participants

The Conditions Affecting Neurocognitive Development and Learning in Early Childhood (CANDLE) study is a U.S.-based observational and longitudinal birth cohort(30). A total of 1503 eligible pregnant women were recruited from Shelby County, Tennessee between 2006 and 2011. The CANDLE study was approved by review boards at the University of Tennessee Health Science Center (UTHSC) and three hospitals where eligible participants could deliver (06-08495-FB). This study uses a subset (N=515) of CANDLE participants with complete DNAm, genotyping, prenatal NO_2_, PM_2.5_, and PM_10_ exposure estimates, and covariate data and was approved by the University of Manitoba Bannatyne Research Ethics Board (HS23413). Covariates were selected *a priori* for inclusion in linear models to account for biological variation and included maternal prenatal tobacco use, adjusted household income, biological sex, gestational age, genetic ancestry, and estimated cell types. See ***SI Methods*** for a list of inclusion/exclusion criteria, information on genotyping preprocessing, quality control, and generation of genetic ancestry principal components (PCs), cell type estimations and PCs, and covariate descriptions.

### Prenatal air pollution exposure estimates

This study examined the separate effects of estimated prenatal exposure to three air pollution components: NO_2_, PM_2.5_, and PM_10_. Prenatal NO_2_ and PM_2.5_ exposure estimates were calculated using a spatiotemporal extension of the universal kriging model at the individual address level that incorporated monitoring data from local monitors and regulatory networks, and land-use characteristics(31, 32). Average pregnancy period NO_2_ and PM_2.5_ exposures were obtained by averaging region-specific biweekly estimates over the pregnancy period (date of conception to date of birth). Prenatal PM_10_ exposure estimates were calculated in a similar manner(33). However, PM_10_ models used a fixed year (2006) for all estimates to avoid temporal confounding or bias related to year-specific modelling errors. Address changes were accounted for using a time-weighted exposure average of reported addresses.

### DNA methylation measurement, preprocessing, and quality control

Umbilical cord blood was collected at birth and processed to obtained cord blood mononuclear cells (CBMCs), which were stored at -80C until use. Genomic DNA was purified using the Qiagen Blood and Tissue Kit (Qiagen, USA), and bisulfite-converted using the EZ-96 DNA Methylation kit (Zymo Research, Irvine, CA, and subsequently hybridized to the Illumina MethylationEPIC array. All nucleic acid procedures followed manufacturer protocol. Multiple R (v3.5) packages including *ewastools*(*34, 35*), *minfi*(*36*), and *conumee*(*37*) were used to perform sample quality checks quality control checks. Briefly, these tools were applied to assess technical parameters, to determine concordance between reported and estimated biological sex, to confirm clustering of samples based on the median intensities in both the methylated and unmethylated channels, to check maternal blood contamination(38), to assess sample identify using SNP probes and to remove samples that have bad detection p-values in >1% of probes, and <3 beads contributing to DNAm signal in >1% of probes. Afterwards, DNAm data were corrected for background noise and normalized for dye-bias using *preprocessNoob* from the *minfi* R package(36), followed by Beta-Mixture Quantile (BMIQ) normalization(39) from the *watermelon* R package(40) to adjust for type I-type II probe bias(36, 40). Poorly performing probes (detection p-value >0.01), probes with <3 beads in >5% of samples, probes on sex chromosomes, cross-hybridizing probes, polymorphic probes, and missing probes were removed, leaving 782,881 probes(41). Lastly, *combat* from the *sva* R package(42) was used to remove batch effects related to chip, chip position, and plate.

### Age four to six years health outcomes

This study investigated the role of air-pollution induced DNAm alterations in wheeze phenotypes. Wheeze phenotypes at ages four to six years were created based on findings from the Tucson Children’s Respiratory Study and included: never, transient, persistent, and late-onset wheeze(14). Participants with never wheeze were defined as those with no reports of wheeze at the year three or year four-six visit. Given that atopic dermatitis and childhood wheeze and asthma are thought to have a shared etiology(43), we also excluded participants that reported atopic dermatitis from the never wheeze group. See ***SI Methods*** for definitions of wheeze used in this study

### Maternal diet

Mothers completed the Block Food Frequency Questionnaire (FFQ) at intake (16 to 26 weeks gestation) and responses were processed by NutritionQuest (Berkley, California) to quantify micronutrient intake, serving size, and frequency of food item consumption. This information was subsequently used to calculate Alternative Healthy Eating Index for Pregnancy (AHEI-P) scores(29). Participants reporting very low (<1000 kcal/day) or very high (>5000 kcal/day) daily energy intake were excluded from further analysis(44).

### Statistical analysis

All statistical analyses were performed using R (v4.2-4.5).

#### Individual and regional DNAm changes

Epigenome-wide analyses were conducted using robust multivariable linear regressions. DMRs were identified by spatially-adjusting p-values obtained from epigenome-wide analyses using the *combp* function(45) from the *missMethyl* R package with seed equal to 0.01. DMRs were excluded from further analysis if they contained fewer than 3 CpGs or if composite CpGs within the DMR exhibited inconsistent directionality in effect sizes.

#### Association with health outcomes

Casual mediation analysis was performed using the *mediation* R package(46) with a quasi-Bayesian approach and 1000 iterations. Health outcomes were modelled as 0 (control) or 1 (affected) in multivariable linear models examining the total effect of prenatal air pollution exposure and treatment. Multivariable linear models also examined the effect of prenatal air pollution exposure on DNAm in mediation analysis. Treatment and control values were defined as mean ± standard deviation of air pollution exposure at the study population level.

#### Buffering by maternal diet micronutrients and overall quality

Robust linear multivariable regressions investigated buffering of DNAm differences. Specifically, an interaction term was included between each air pollutant, respectively, and vitamin C, the ratio of n-6 to n-3 PUFA consumption, folate, or AHEI-P score. We considered three potential indicators of buffering by diet: (1) Re-identification of DMRs from the main analysis within the interaction terms of the diet-adjusted models, suggesting these DNAm differences may depend on maternal nutrient intake or overall diet quality; (2) The emergence of new DMRs in the main air pollutant term of the diet interaction models, which may indicate DNAm differences that were previously masked by maternal diet in the primary analysis; (3) New or stronger causal mediation effects for DMRs identified in the main analysis that emerged only after including an interaction term between air pollutants and maternal intake of vitamin C, PUFAs, folate, or AHEI-P score (47). For DMRs exhibiting new or stronger causal mediation effects, we attempted to quantify the significance of buffering using the *mediation* R package(46) to conduct moderated mediation. Moderated mediation models were specified as described above, with the addition of an interaction term between air pollutant exposures and individual micronutrients or AHEI-P score. Casual mediation analysis was performed using the *mediation* R package(83) with a quasi-Bayesian approach and 10 iterations. Mediator treatment and control values were defined as mean ± standard deviation of air pollution exposure at the study population level. Moderated was tested using the *test.modmed()* function from the mediation R package with a quasi-Bayesian approach and 1000 iterations. Moderator levels were defined as mean ± standard deviation of micronutrients or AHEI-P score.

#### Statistical considerations

The MASS R package(48) was used to run multivariable linear regressions with the Huber’s Proposal-2 M-estimator. For all analyses (unless otherwise specified), models were run twice: once with DNAm m-values to obtain p-values, as m-values are homoscedastic and therefore more statistically appropriate, and again with β-values to obtain biologically-interpretable effect sizes(49). CpGs that failed to converge within 50 iterations were removed from further analysis in epigenome-wide investigations. All models included variables to adjust for biological variation and technical variation (see ***SI Methods***). Vitamin C, PUFA, and folate intake and AHEI-P scores were mean-centred prior to inclusion in robust linear models. P-values obtained from epigenome-wide investigations were adjusted using the Benjamini-Hochberg method. P-values obtained from (moderated) mediation analyses were adjusted using the Bonferroni method. Air pollutant effect sizes (β-values) were defined as the change in DNAm across the range of air pollution exposure estimates and were calculated by multiplying the coefficient of air pollution exposures obtained from robust multivariable linear models by the range of air pollution exposures in the study population. Effect sizes of micronutrients or overall diet quality buffering on air pollutant-associated DNAm differences (β-value/ppb or β-value/ug/m^3^) were defined as the change in DNAm across the range of micronutrient or diet quality exposure and were calculated by multiplying the coefficient of the interaction term between a given micronutrient or overall diet quality obtained from robust multivariable linear models by the range of micronutrient intake or maternal diet quality across. Statistical significance was defined as (adjusted) p<0.05 and for epigenome-wide and biological significance was defined as an absolute effect size >0.01% for individual and region DNAm differences.

## Results

A total of 515 mother-child dyads from the socioeconomically and racially heterogeneous cohort of CANDLE participants (30) were included in our main analyses of prenatal air pollution exposures. Most mothers self-identified as either Black/African American (53%) or White (40%), with the remainder identifying as Asian (1%), American Indian/Alaskan Native (<1%), as multiple races (5%), or another race not listed on the questionnaire (1%; **Table 1**). Median annual prenatal PM_2.5_ and NO_2_ exposure estimates of 10.5 μg/m^3^ and 8.1 ppb were near or below the current primary annual standards (9 μg/m^3^ and 53 ppb, respectively) for human health protection set by the United States Environmental Protection Agency (US EPA)(50, 51); there is no USA EPA annual standard for PM_10_ exposure but we noted a median PM_10_ exposure of 17.7 ppb(52). A total of 13%, 10%, and 4% of study participants exhibited transient, persistent, or late-onset wheeze, respectively, with 12% of biological males and 8% of female exhibiting persistent wheeze (**Table S1**). Caregiver reports of child atopic dermatitis in the study population were high (25%) compared to U.S. prevalence rates of 13.7% in children age zero to five years old(53), and even higher in those with transient (43%), persistent (45%), or late-onset (37%) wheeze (**Table 1 and Table S2**). Only 6% of those with transient wheeze received an asthma diagnosis by the age four to six year visit, compared to 82% of individuals with persistent wheeze and 58% with late-onset wheeze (**Table S2**). Participants with transient, persistent, or late-onset wheeze also exhibited consistent trends of higher prenatal exposure to NO_2_, PM_2.5_, and PM_10_ compared to individuals without wheeze, though differences did not reach statistical significance (one-sided t-test p>0.05; **Fig S1**).

**Table 1.**
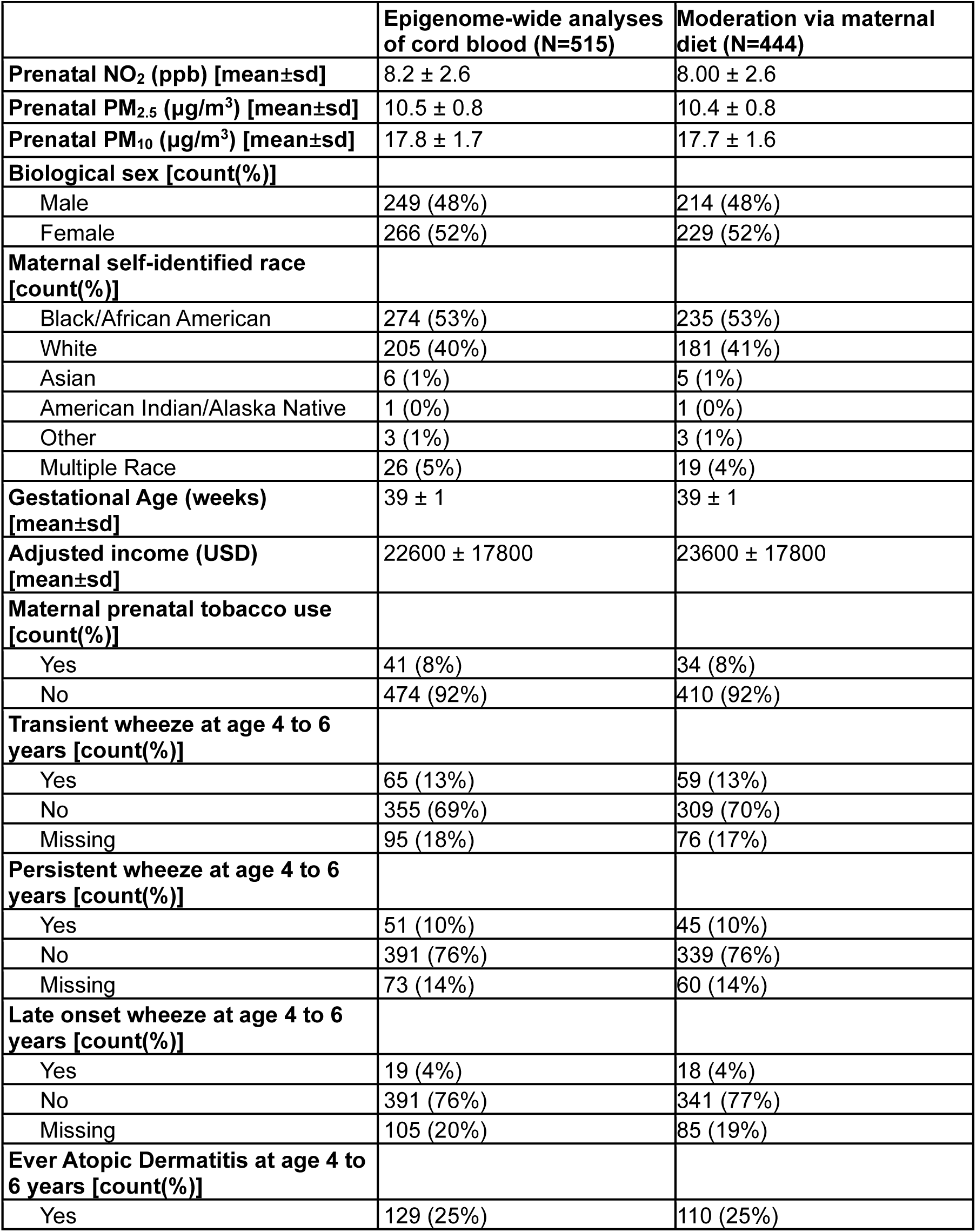
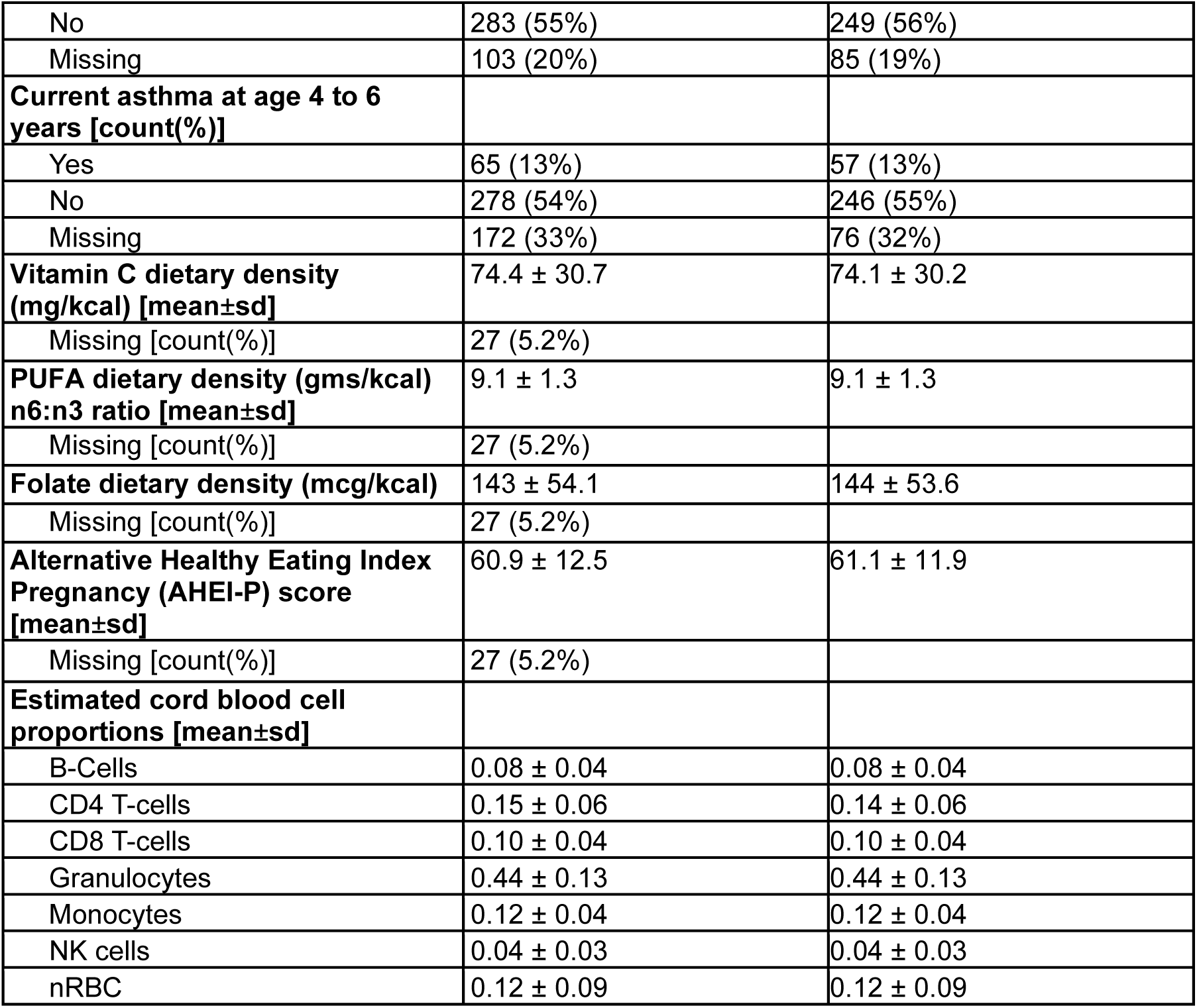
Study population characteristics of mother-child dyads included in the main epigenome-wide prenatal air pollutant exposures (N=515) and in the secondary analysis examining moderation by maternal prenatal diet factors (N=444) analysis sets.

|  | Epigenome-wide analyses of cord blood (N=515) | Moderation via maternal diet (N=444) |
| --- | --- | --- |
| **Prenatal NO<sub>2</sub> (ppb) [mean±sd]** | 8.2 ± 2.6 | 8.00 ± 2.6 |
| **Prenatal PM<sub>2.5</sub> (µg/m<sup>3</sup>) [mean±sd]** | 10.5 ± 0.8 | 10.4 ± 0.8 |
| **Prenatal PM<sub>10</sub> (µg/m<sup>3</sup>) [mean±sd]** | 17.8 ± 1.7 | 17.7 ± 1.6 |
| **Biological sex [count(%)]** |  |  |
| Male | 249 (48%) | 214 (48%) |
| Female | 266 (52%) | 229 (52%) |
| **Maternal self-identified race [count(%)]** |  |  |
| Black/African American | 274 (53%) | 235 (53%) |
| White | 205 (40%) | 181 (41%) |
| Asian | 6 (1%) | 5 (1%) |
| American Indian/Alaska Native | 1 (0%) | 1 (0%) |
| Other | 3 (1%) | 3 (1%) |
| Multiple Race | 26 (5%) | 19 (4%) |
| **Gestational Age (weeks) [mean±sd]** | 39 ± 1 | 39 ± 1 |
| **Adjusted income (USD) [mean±sd]** | 22600 ± 17800 | 23600 ± 17800 |
| **Maternal prenatal tobacco use [count(%)]** |  |  |
| Yes | 41 (8%) | 34 (8%) |
| No | 474 (92%) | 410 (92%) |
| **Transient wheeze at age 4 to 6 years [count(%)]** |  |  |
| Yes | 65 (13%) | 59 (13%) |
| No | 355 (69%) | 309 (70%) |
| Missing | 95 (18%) | 76 (17%) |
| **Persistent wheeze at age 4 to 6 years [count(%)]** |  |  |
| Yes | 51 (10%) | 45 (10%) |
| No | 391 (76%) | 339 (76%) |
| Missing | 73 (14%) | 60 (14%) |
| **Late onset wheeze at age 4 to 6 years [count(%)]** |  |  |
| Yes | 19 (4%) | 18 (4%) |
| No | 391 (76%) | 341 (77%) |
| Missing | 105 (20%) | 85 (19%) |
| **Ever Atopic Dermatitis at age 4 to 6 years [count(%)]** |  |  |
| Yes | 129 (25%) | 110 (25%) |
| No | 283 (55%) | 249 (56%) |
| Missing | 103 (20%) | 85 (19%) |
| **Current asthma at age 4 to 6 years [count(%)]** |  |  |
| Yes | 65 (13%) | 57 (13%) |
| No | 278 (54%) | 246 (55%) |
| Missing | 172 (33%) | 76 (32%) |
| **Vitamin C dietary density (mg/kcal) [mean±sd]** | 74.4 ± 30.7 | 74.1 ± 30.2 |
| Missing [count(%)] | 27 (5.2%) |  |
| **PUFA dietary density (gms/kcal) n6:n3 ratio [mean±sd]** | 9.1 ± 1.3 | 9.1 ± 1.3 |
| Missing [count(%)] | 27 (5.2%) |  |
| **Folate dietary density (mcg/kcal)** | 143 ± 54.1 | 144 ± 53.6 |
| Missing [count(%)] | 27 (5.2%) |  |
| **Alternative Healthy Eating Index Pregnancy (AHEI-P) score [mean±sd]** | 60.9 ± 12.5 | 61.1 ± 11.9 |
| Missing [count(%)] | 27 (5.2%) |  |
| **Estimated cord blood cell proportions [mean±sd]** |  |  |
| B-Cells | 0.08 ± 0.04 | 0.08 ± 0.04 |
| CD4 T-cells | 0.15 ± 0.06 | 0.14 ± 0.06 |
| CD8 T-cells | 0.10 ± 0.04 | 0.10 ± 0.04 |
| Granulocytes | 0.44 ± 0.13 | 0.44 ± 0.13 |
| Monocytes | 0.12 ± 0.04 | 0.12 ± 0.04 |
| NK cells | 0.04 ± 0.03 | 0.04 ± 0.03 |
| nRBC | 0.12 ± 0.09 | 0.12 ± 0.09 |

The main analyses of this study examined associations between prenatal air pollutant exposures and cord blood DNAm among the 515 CANDLE participants. Specifically, we conducted epigenome-wide analyses of NO_2_, PM_2.5_, and PM_10_ to identify potential novel DNAm associations with broader applicability across human populations, and to assess whether findings previously reported in predominantly European ancestry populations replicate in a more diverse study population (**Fig S2**; **Files S1-3**). Models exhibited good genomic inflation factors between 0.93 and 1.04 (54). A single CpG (cg15994137) annotated to the body of *ZFYVE9* was significantly and negatively associated with prenatal PM_2.5_ exposure (adjusted p=0.046, effect size=-0.06). No other statistically significant cord blood DNAm differences were identified in the epigenome-wide analyses of either prenatal NO_2_ or prenatal PM_10_ exposure.

As part of our main analyses, we next examined differentially methylated regions (DMRs)(45), which provide greater power to detect small DNAm differences compared to epigenome-wide investigations. Prenatal exposures to NO_2_, PM_2.5_, and PM_10_ were associated with 19, seven, and five cord blood DMRs, respectively (Sidak p<0.05 and mean absolute effect size>0.01; **Table 2**). Although no DMRs were shared between air pollutants (**Fig 1**), each air pollutant was associated with at least one DMR annotated to an immune-relevant gene including members of the *HLA* family (NO_2_ and PM_2.5_)(55), *IL9* (PM_2.5_)(56), and *GCNT2* (PM_10_)(57). We further examined whether any of the observed regional DNAm differences might be under the influence of nearby genetic variations by crosschecking CpGs within each DMR with an online mQTL database (58). At least half of the component CpGs in three NO_2_-associated DMRs, one PM_2.5_-associate DMR, and one PM_10_-associated DMR were previously annotated as mQTLs, suggesting that DNAm differences in these regions may be influenced by nearby genetic variants (**Table 2**). Allelic effect sizes (β-value) of potential mQTLs identified in the analyses of NO_2_, PM_2.5_, and PM_10_ exposure ranged from 0.02 to 0.28, 0.02 to 0.16, and 0.02 to 0.8, respectively.

**Table 2.**
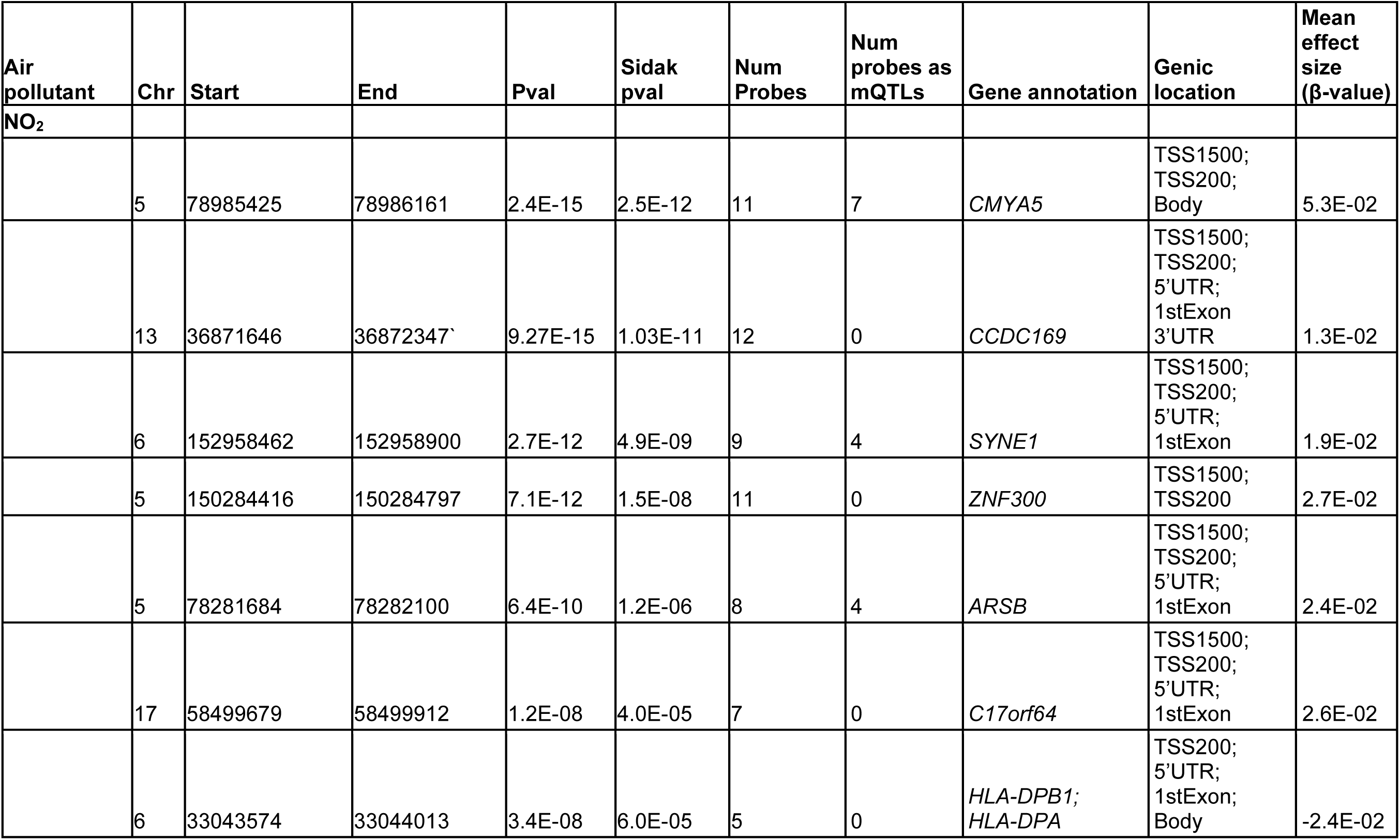
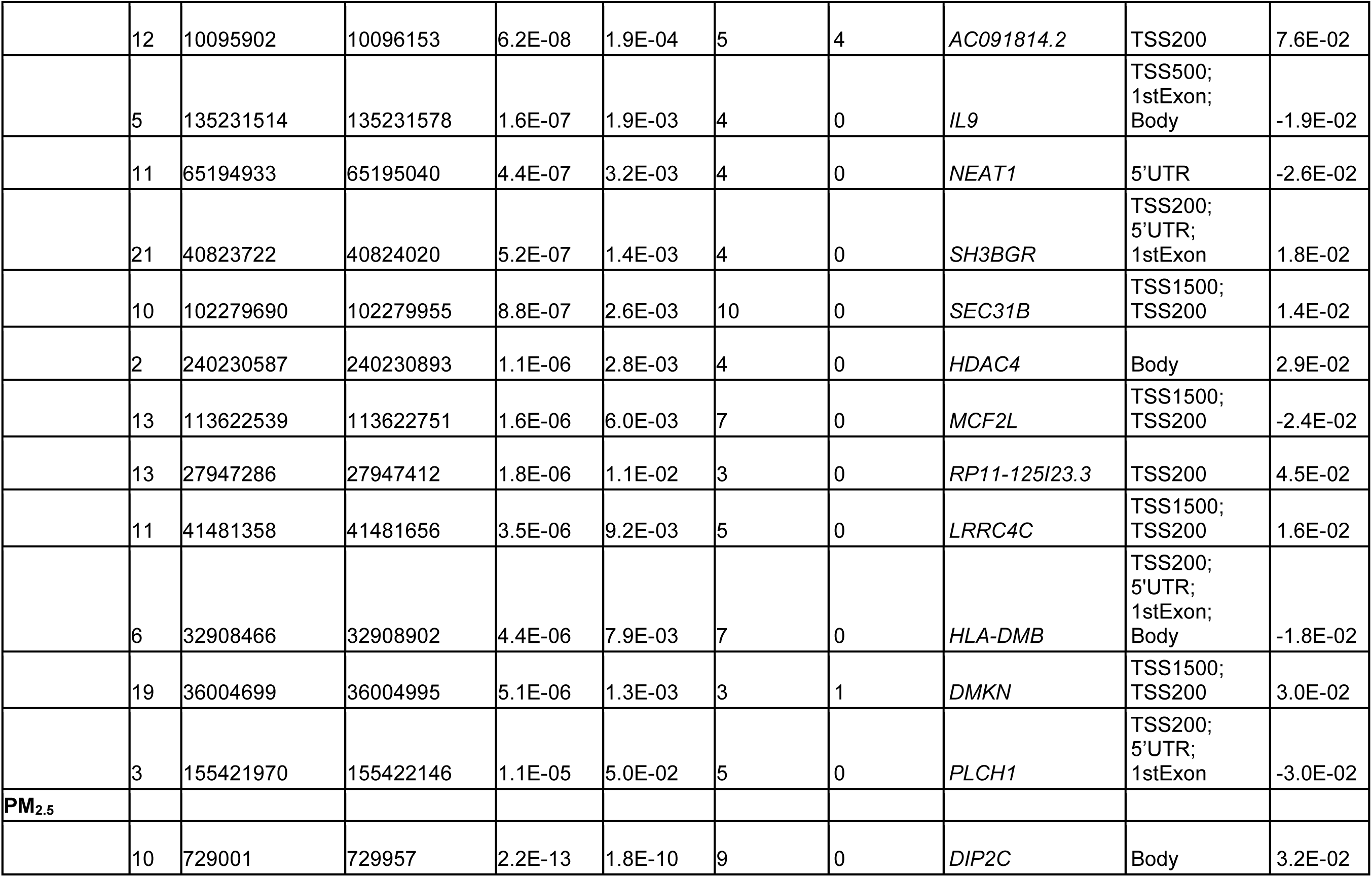
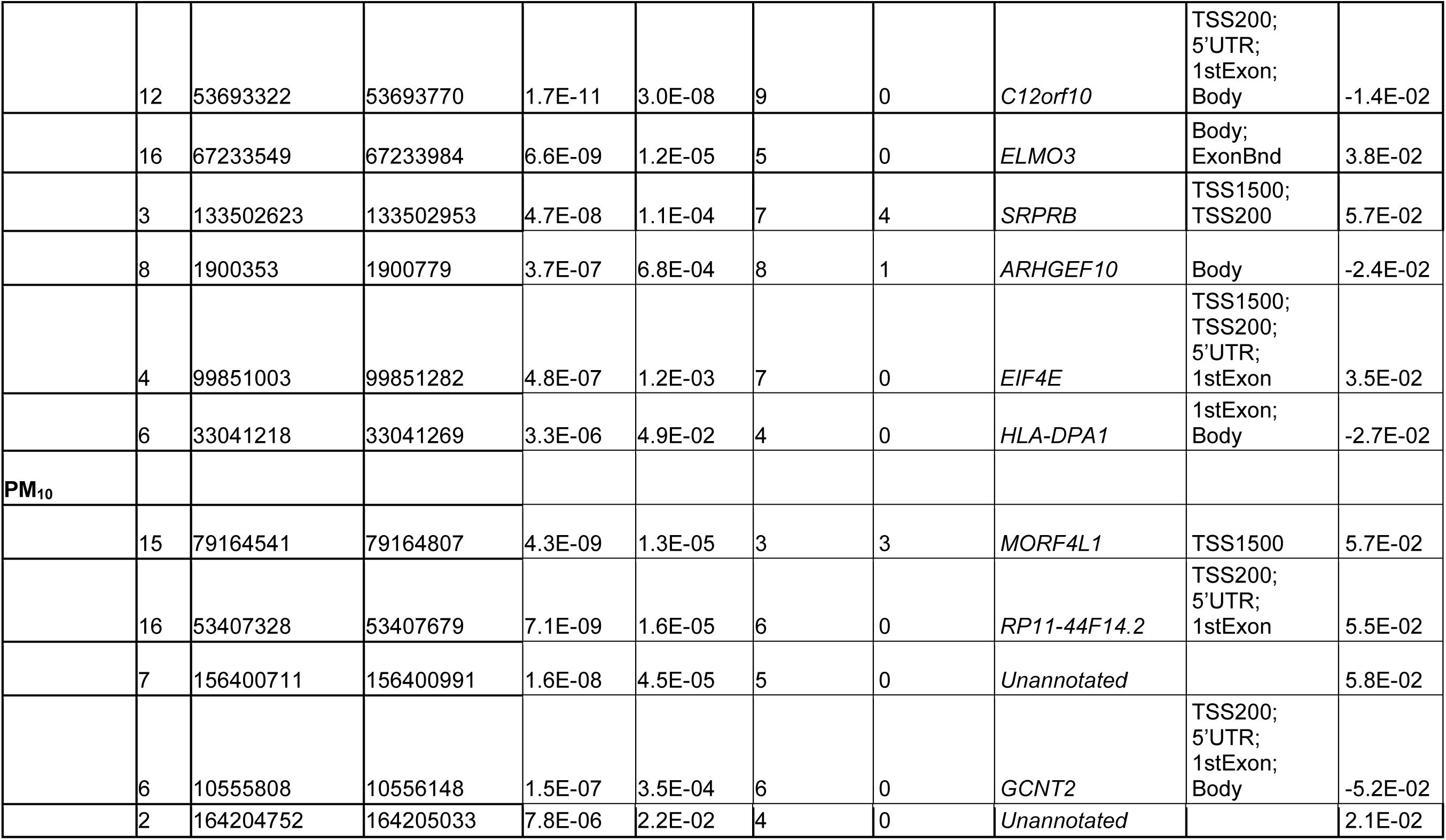
Differentially methylated regions in cord blood that are associated with prenatal NO_2_ (N=19), PM_2.5_ (N=7), and PM_10_ (N=5) exposures in CANDLE participants (N=515).

**Figure 1.**
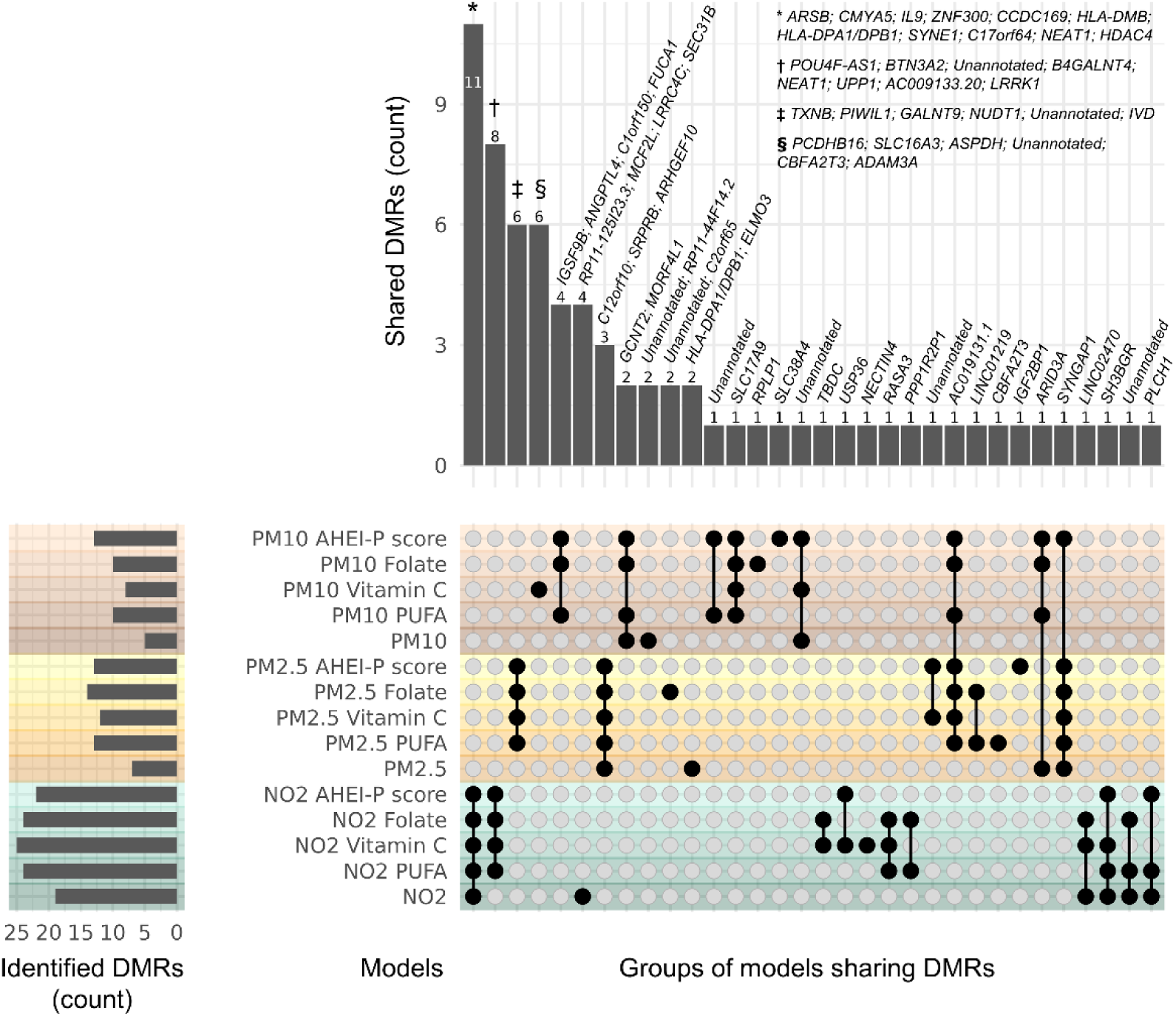
Overlap of differentially methylated regions (DMRs) identified in the main air pollution exposure analysis (N=515 participants) or in the main effect term of NO_2_, PM_2.5_, or PM_10_ in secondary diet interaction analyses (N=444 participants). Diet interaction analysis account for the maternal intake of vitamin C, the ratio of n-6 to n-3 polyunsaturated fatty acids (PUFAs), or overall diet quality using Alternative Healthy Eating Index Pregnancy (AHEI-P) scores. The bar chart on the left of the graph displays the total number of DMRs identified in each analysis. The dot matrix indicates which groups of DMRs belong to which analysis. An analysis that contains a group of DMRs is specified by a black circle. Black vertical lines joining plots indicate that a group of DMRs was identified in one or more analysis. The bar chart on the top of the graph indicates the number of DMRs in a given group. Bars are annotated with the genic annotations of DMRs within that group.

The biological embedding of prenatal air pollution exposures via DNAm is hypothesized to contribute to child respiratory disease including wheeze and asthma(24). Although we consistently observed wheeze phenotypes were positively associated with air pollutant exposures (except for adjusted models of NO_2_ and persistent wheeze, and PM_2.5_ or PM_10_ and late-onset wheeze), the difference in prenatal air pollution exposures between non-wheezing and wheezing individuals did not reach statistical significance (defined as p<0.05) in one-sided t-tests (**Fig S1**). Nevertheless, we proceeded to investigate the role of mean DNAm across DMRs associated with prenatal NO_2_, PM_2.5_, and PM_10_ in wheeze outcomes using causal mediation analysis (**Files S4-S12**). One DMR within the promoter of chr6:33043574–33044013 (*HLA-DPA1/DPB1*) showed a suggestive positive mediation effect on transient wheeze associated with prenatal NO_2_ (unadjusted p=0.032, ACME=0.040; **Fig 2A**). A nearby DMR in this same region (chr6:33041218-33041269) also showed a suggestive mediation effect on transient wheeze in relation to PM_2.5_ exposure (unadjusted p=0.038, ACME=4.7E-02; **Fig 2B**). In both cases, the average direct effects of air pollution (-4.4E-02, and -8.2E-03, respectively) were in the opposite direction of the mediation effects. However, positive mediation effects did not remain significant after multiple testing correction (**Files S4-12**).

**Fig 2.**
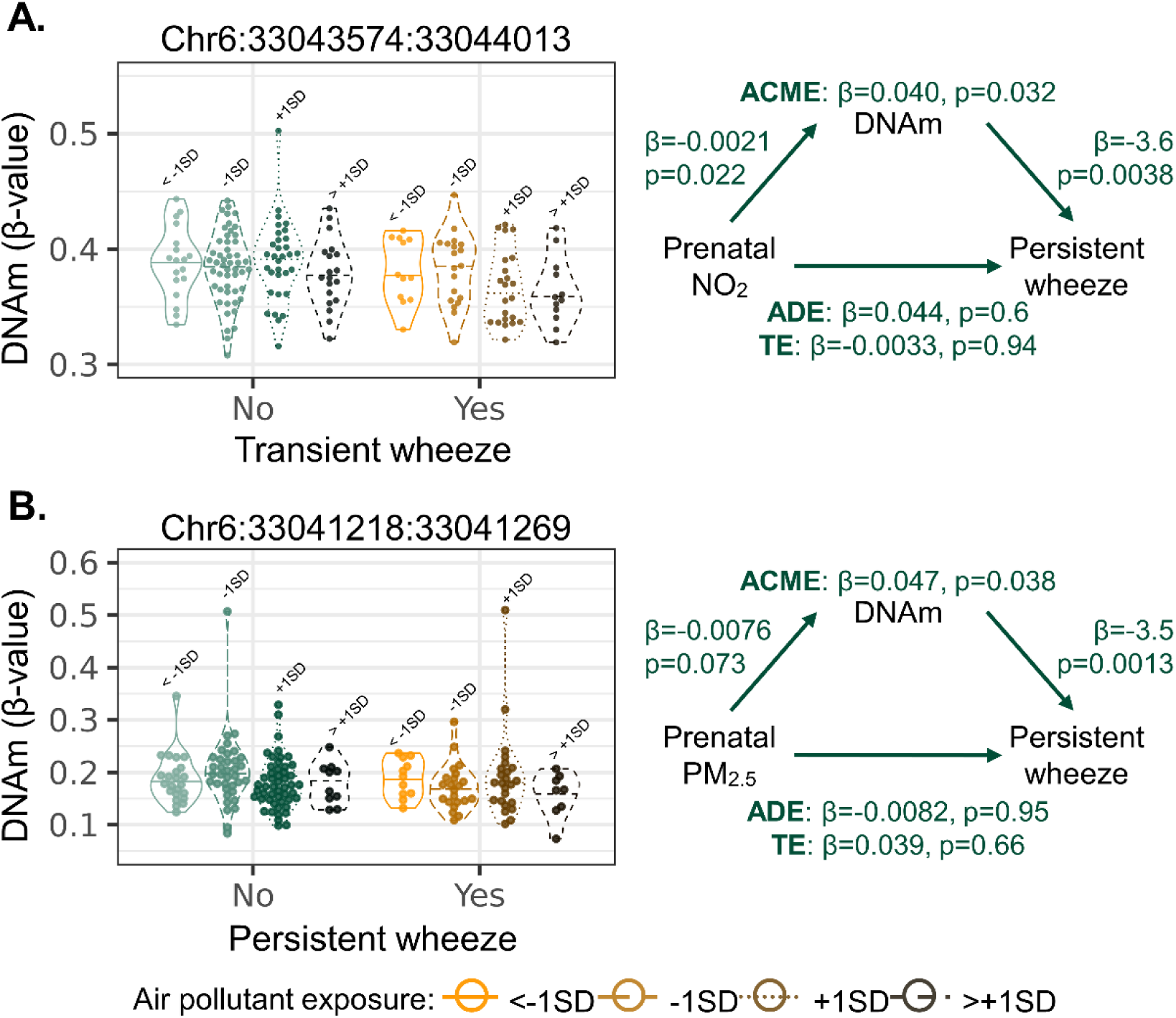
Causal mediation effects of cord blood differentially methylated regions (DMRs) between prenatal NO_2_ or PM_2.5_ exposures and transient wheeze (N=65 with, 119 without). (**A**) Relationship between mean DNAm of chr6:33043574-33044013 (*HLA-DPA1/DPB1*), prenatal NO_2_ exposure, and transient wheeze (*left*). Mediation pathway of prenatal NO_2_ exposure on transient wheeze via mean chr6:33043574-33044013 DNAm (*right*). (**B**) Relationship between mean DNAm of chr6:33041218-33041269 (*HLA-DPA1*), prenatal PM_2.5_ exposure, and transient wheeze (*left*). Mediation pathway of prenatal PM_2.5_ exposure on transient wheeze via mean chr6:33041218-33041269 DNAm (*right*). ACME = average causal mediation effects; ADE = average direct effects; TE = total effects. Participants with transient wheeze (N=65) are coloured in shades of orange/brown, and participants without transient wheeze (N=119) coloured in shades of green, where darker shades represent higher air pollution exposures. Data are stratified by NO_2_ exposure quartiles for visualization. The first quarter is represented by open circles and solid lines, the second quarter by open triangles and long dashed lines, the third quarter by open squares and dotted lines, and the fourth quarter by crosses and short dashed lines.

We next tested the hypothesis that maternal intake of protective micronutrients including vitamin C(27), or the ratio of n6 to n3 PUFAs (59, 60), or folate(26) could buffer the effects of prenatal exposure to NO_2_, PM_2.5_, or PM_10_ on DMRs identified in our main analyses(35). Because these beneficial micronutrients were correlated to one another (**Fig S3**), we also examined the potential buffering effect of overall diet quality using the AHEI-P score. This score ranges from 0 to 130, with higher values indicating greater intake of nutrients associated with chronic disease prevention(61). To test the above hypothesis, we conducted a secondary analysis, repeating epigenome-wide analyses (**Files S13-S24**), but this time including interaction terms between each air pollutant (NO_2_, PM_2.5_, PM_10_) and either each micronutrient or the AHEI-P score. We focused on DMRs from our main analysis that had both a significant pollutant main effect term and a significant interaction term of the new models (**Files S25-S45**), with effect sizes in opposite directions, indicating a potential buffering effect. Only one NO_2_-associated DMR annotated to *C17orf64* (chr17:58499679-58499912) from our main analysis met this criterion when assessing the effect of maternal folate intake in secondary analyses. Specifically, higher maternal folate intake was associated with lower DNAm across this DMR (effect size = -1.23E-02 β-value/ppb), counteracting the higher DNAm linked to NO_2_ exposure (**Fig 3A**). Though neither individual micronutrients nor overall diet quality significantly buffered other DMRs identified in the main analyses, we observed directionally consistent buffering effects for one or more micronutrients or for AHEI-P across 16 of the 19 NO₂-associated DMRs and across all PM_2.5_- and PM_10_-associated DMRs(**Fig 3B-D; Files 46-57**). Absolute interaction effect sizes were biologically significant (≥1.00×10⁻² β-value/ppb or β-value/μg/m³) for two NO₂-associated DMRs, five PM_2.5_-associated DMRs, and four PM_10_-associated DMRs, suggesting buffering may be occurring across these DMRs even though they were not identified in any of the interaction effect terms.

**Fig 3.**
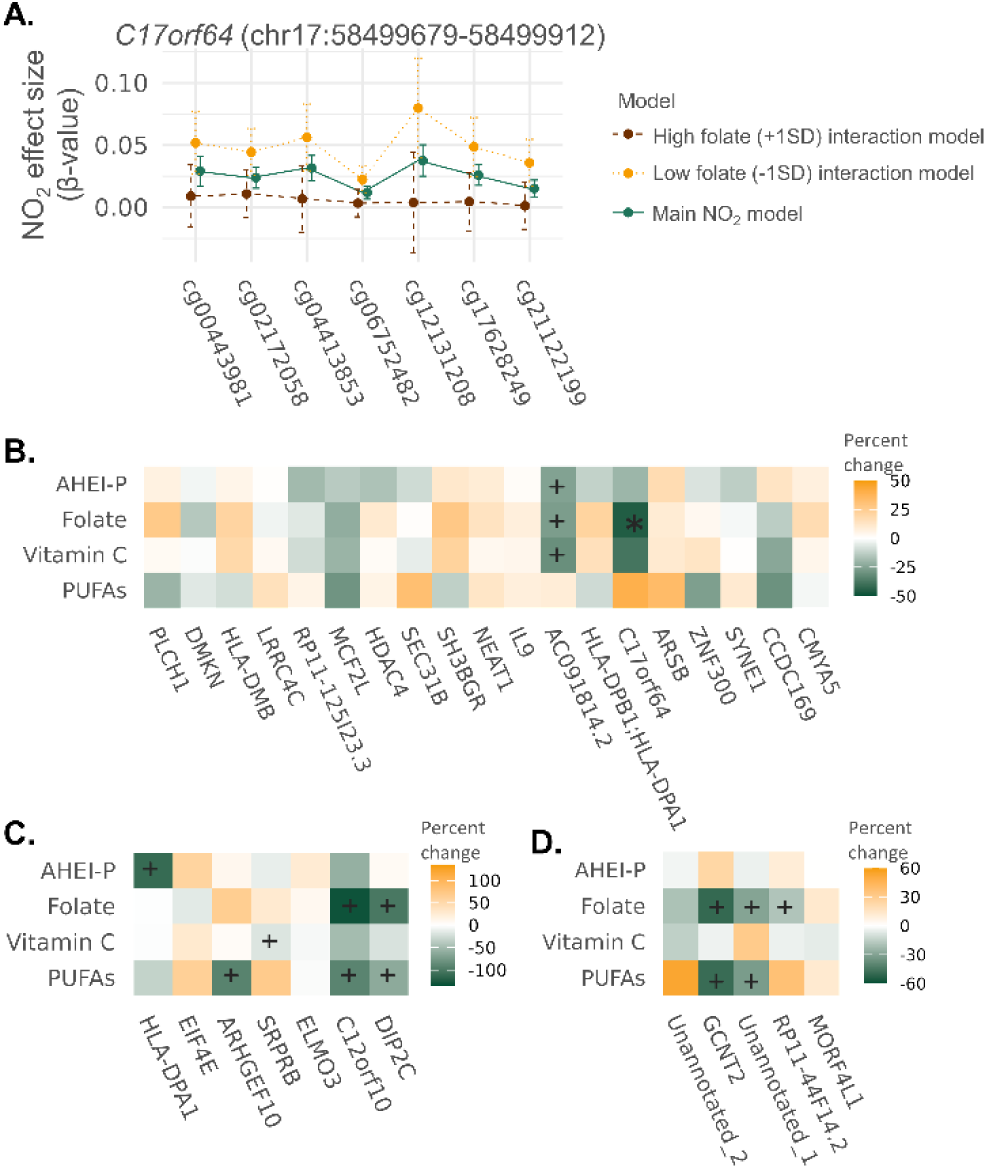
Effect sizes of the interaction term between maternal intake of polyunsaturates fatty acids (PUFAs), vitamin C, folate, or Alternative Healthy Eating Index -Pregnancy (AHEI-P) and air pollutants from secondary analyses relative to the main effect of NO_2_, PM_2.5_, or PM_10_ across cord blood differentially methylated regions (DMRs) identified in the main analyses of CANDLE participants (N=515). (**A**) A DMR associated with NO_2_ and annotated to *C17orf64* in the main analysis was also identified in the interaction term of the model accounting for maternal folate intake, indicating a potential buffering effect. Across the DMR, mean DNAm associated with NO_2_ was lower when accounting for higher maternal folate intake (+1SD) and increased when considering low maternal folate intake (-1SD). In interaction models, maternal intake of PUFAs, vitamin C, folate, and overall diet quality (AHEI-P) reduced the effect size of (**B**) NO_2_, (**C**) PM_2.5_, and (**D**) PM10 across several of the DMRs from the main analysis. Buffering effect sizes are displayed in percent (%) change of interaction effect sizes relative to the main effect sizes of air pollutants. Buffering is considered when the percent change is negative, representing an interaction effect in the opposite direction of the main air pollutant effect. A positive percent change occurs when the interaction effect and main effect are in the same direction and is not considered beneficial. Plus (+) symbols indicate interaction effect sizes that surpassing biological significance (≥0.01 β-value/ppb or β-value/μg/m³) but not identified in the interaction effect term of any of the interaction models. The asterisk (*) symbol indicates the NO_2_ associated DMR annotated to *C17orf64* that was identified in both the main effect term and interaction effect term.

We were also interested in DMRs that emerged in the air pollutant main effect term of the interaction models but were not identified in our main analyses (**Files S24-S36**). These DMRs likely represent pollutant-associated DNAm changes that were previously masked by the modifying effects of maternal micronutrient intake and/or overall diet quality. More than half of the DMRs associated with the main effect of PM_2.5_ or PM_10_ in diet interaction models (vitamin C, PUFAs, folate, and AHEI-P) were not detected in the main analyses (**Fig 1**). Similarly, over a third of DMRs associated with the main effect of NO_2_ in diet interaction models were not detected in main analyses. Among these, a NO_2_-associated DMR annotated to *UPP1* (chr7:48129797-48130198) exhibited positive causal mediation effects (0.043 to 0.046) on the relationship between NO_2_ exposure and persistent wheeze in diet interaction models that included PUFAs, folate, or AHEI-P score (**Files S58-S60**). These mediation effects were in the opposite direction of the direct effect of NO_2_ on persistent wheeze and did not remain significant after Bonferroni correction (unadjusted p=0.024 to 0.032). No other newly emerged DMRs had significant positive causal mediation effects between any air pollutant and wheeze phenotype (**Files S61-S93**). Most (N=8 of 12) newly emerged NO₂-associated DMRs were shared across all diet interaction models (**Fig 1**), likely reflecting the high correlation between micronutrients (**Fig S4**). A similar pattern was observed for PM_2.5_ DMRs where 7 of 9 newly emerged DMRs were shared across diet interaction models. In contrast, most newly identified PM_10_-associated DMRS were shared only across diet interaction models adjusting for PUFA intake, folate, or AHEI-P score, but not vitamin C (**Fig 1**), which suggests maternal intake of vitamin C may have a less prominent role in buffering the effects of PM_10_ on cord blood DNAm.

Most NO_2_ (≥13 of 19) and PM_2.5_ (4 of 7) DMRs identified in our main analysis were also identified in the air pollutant effect term across all diet interaction models (**Fig 1**; **File S24-S30**). When we repeated mediation analyses with interaction terms to account for maternal dietary effects, we identified new positive causal mediation effects between NO_2_ and transient wheeze through a DMR annotated to *HLA-DMB* in models adjusting for vitamin C, folate, or AHEI-P score (**Files S61**, **S63**, and **S64**). This DMR also showed positive mediation effect between NO_2_ and late wheeze when adjust for vitamin C, PUFAs, folate, or AHEI-P score (**Files S66-S69**). A second DMR annotated to *SYNE1* exhibited positive mediation effects between NO_2_ and transient wheeze only after adjusting for vitamin C or PUFA intake (**Files S61** and **S62**). Additionally, we observed that the positive causal mediation effects of *HLA-DPA1/DPB1* on the relationship between NO₂ and transient wheeze increased from 0.04 to as much as 0.06 after adjustment for intake of vitamin C, PUFA, or folate or AHEI-P score (**Files S61-S64**). Although none of these causal mediation effects reached statistical significance after multiple testing correction, the emergence or amplification of these effects following dietary adjustment(47) supports our hypothesis that maternal intake of beneficial micronutrients and overall diet quality may buffer the effects of air pollutants observed in the main models. We further evaluated this hypothesis using moderated mediation analysis; however, the difference in causal mediation effects between ±1 standard deviation of vitamin C, PUFAs, folate, and AHEI-P did not reach statistical significance for any of the DMRs described above (**File S94-S98**).

Previous research has shown that biological males are at greater risk of developing childhood asthma following prenatal exposure to air pollution(11–13). However, the role of sex-specific DNAm changes in this relationship remains unclear. To investigate this, we repeated the epigenome-wide analyses, this time including an interaction term between each air pollutant and biological sex (**File S99-101)**. Our primary focus was on DMRs identified in the interaction term, as these regions represent locations where the effects of air pollution differ between males and females. The effects of NO_2_, PM_2.5_, and PM_10_ on DNAm differed by biological sex across two, 11, and two DMRs respectively (**File S102-S104**). Males exhibited higher DNAm at both NO_2_-associated DMRs and lower DNAm at six of the 11 PM_2.5_- and both PM_10_-associated DMRs compared to females. At least half of the component CpGs in the six PM_2.5_-associated DMRs were previously annotated as putative mQLTs, with effect sizes ranging from 0.02 to 0.11, indicating potential genetic influence on DNAm across these regions. Several of the PM_2.5_-associated DMRs containing putative mQTLs, including *SERPINB9*(62) *and WFDC1*(63), have previously been linked to lung function in genome- or epigenome-wide studies. However, in this study, none of the sex-differing DMRs showed evidence of mediating the relationships between air pollutants and transient, persistent, or late-onset wheeze. We did not examine mediation effects of sex-differing DNAm regions on late onset wheeze due to small sample size (N=9 males, 10 females).

Finally, we assessed consistency with prior epigenome-wide studies by examining whether CpGs previously associated with prenatal NO_2_, PM_2.5_, or PM_10_ exposure exhibited the same direction of effect in our epigenome-wide investigations, even if they did not reach statistical significance (FDR <0.05). Several DNAm alterations associated with prenatal PM_2.5_ and PM_10_ exposure in Precision Medicine Intervention in Severe Asthma (PRISM) participants, who have similar demographics to CANDLE, were replicated in this study, exhibiting the same direction of effect. These included lower DNAm with greater PM_2.5_ exposure at cg01011943 (*PSG5*) and cg00348551 (*C7orf50*), and higher DNAm with PM_10_ exposure at cg00905156 (*FAM13A*) and cg06849931 (*NOTCH4*) (**Fig S15A-E**) (19, 20). We also observed higher DNAm at cg18684142 (*HOXB6*) with NO_2_ in CANDLE participants, consistent with findings from the primarily European CHILD cohort study(20), implying this change is robust across populations. In contrast, several findings from studies using primarily European cohorts did not replicate in CANDLE. These include higher DNAm at cg10430077 (*TRIM41*) in males with NO_2_(20), higher DNAm at cg00035623 (*UBE2S*) with PM_2.5_(18), higher DNAm at cg02015529 (*TM9SF2*) with PM_10_(18), and lower DNAm at cg22329831 (*TDRD6*) with PM_10_(18), suggesting these associations may not be generalizable across more diverse populations (**Fig S5F-I**).

## Discussion

In this study we identified DNAm alterations in cord blood associated with prenatal exposure to NO_2_, PM_2.5_, and/or PM_10_ in a subset of participants from the socioeconomically and racially diverse CANDLE cohort study. Most identified DMRs were unique to individual air pollutants, suggesting that different cellular pathways may be affected by prenatal exposure to NO_2_, PM_2.5_, and PM_10_. Altered DNAm in *HLA-DPA1/DBP1* DNAm displayed suggestive positive mediation effects between prenatal exposure to NO_2_ and PM_2.5_ and transient wheeze. Additional DMRs associated with prenatal air pollutant exposures were identified only after adjusting for maternal intake of protective nutrients or overall diet quality in our secondary analyses, suggesting that the effects of these exposures on DNAm were likely masked by the buffering influence of maternal diet in the initial models. Two DMRs from the main analysis exhibited significant mediation effects only after considering diet interaction terms in the mediation analysis. Additionally, including diet interaction terms in the mediation analysis strengthened the previously observed mediation at *HLA-DPA1/DPB1*. Together, these findings suggest that prenatal air pollutant exposures result in unique alterations to cord blood DNAm, and that nutrient dense, high quality maternal diets may have a role in buffering some of these DNAm differences.

Lower DNAm in the promoter region of *HLA-DPA1/DPB1* displayed suggestive positive causal mediation effects between exposure to NO_2_ or PM_2.5_ and transient wheeze in our main models. These HLA genes code for MHC class II member proteins found on antigen presenting cells and play roles in immune response following inhalation of heavy metals(64) and in bronchodilator response(63). Genome-wide studies have also identified associations between *HLA-DPA1/DPB1* and atopic dermatitis(65) and asthma(66). The role of lower *HLA-DPA1/DPB1* DNAm in transient wheeze associated with NO_2_ is unclear, but could involve similar mechanisms to its function in asthma, where heightened transcription has been shown to lead to increased T-cell activation and inflammation(67).

A primary focus of this work was whether maternal diet could mitigate the effects of prenatal air pollutant exposure on DNAm changes, and ultimately wheeze and asthma in childhood. Positive causal mediation effects of lower *HLA-DPA1/DPB1* DNAm associated with NO_2_ further increased in diet interaction models (vitamin C, PUFA, folate, and AHEI-P score), suggesting micronutrient rich, high-quality diets may play a role in buffering the effects of NO_2_ on DNAm(47). These observations are consistent with previous findings demonstrating greater maternal diet density of vitamin C can reduce the effects of maternal prenatal smoking on DNA and is protective towards child respiratory health at age five(27, 28), and that higher maternal folate can buffer the effects of prenatal PM_10_ exposure on childhood IQ(26). Though differences in the causal mediation effects of *HLA-DPA1/DPB1* DNAm between low and high maternal micronutrient intake did not reach significance in moderated mediation analyses, the above findings support that more healthful maternal diets can help mitigate infant DNAm changes associated with prenatal air pollutant exposures, which could help reduce changes to children’s immune function that contribute to wheeze and asthma.

This study extends prior work by examining the role of air pollution-induced DNAm differences in specific wheeze phenotypes including no/infrequent wheeze, transient wheeze, persistent wheeze, and late onset wheeze at age 4 to 6 years. We based our classifications on those outlined in the original Tucson Study(14). Our ability to classify wheeze phenotypes in CANDLE participants was facilitated by the availability of repeated questionnaire and clinical visit data from birth until the age four clinic visit. Prenatal air pollution exposures have been associated with transient, persistent, and late-onset wheeze phenotypes; however, evidence is limited and findings vary across studies(15). This work extends our knowledge of prenatal air pollution exposure and respiratory health by providing insight into the air pollution-induced DNAm differences associated with wheeze phenotypes. Specifically, these findings suggest that prenatal NO_2_ and PM_2.5_ may contribute to transient wheeze phenotypes in part through a shared pathway involving HLA-DP genes. HLA-DP genetic variants have been also associated with atopic dermatitis(43), and we noted a high rate of atopic dermatitis in participants with transient wheeze here and in the full study cohort compared to the Tucson Children’s Respiratory Study participants(14). It remains possible that shared genetic variants are driving the development of wheeze and atopic dermatitis in this study, but is unlikely given that we adjusted data for genetic ancestry and HLA-DP gene CpGs did not constitute previously identified mQTLs listed on mQTLdb(58). Replication of these findings in other racially diverse cohorts will help confirm a shared role for the regulation of HLA-DP genes in the relationship between prenatal air pollutant exposures and childhood wheeze and asthma.

Biological males are more susceptible to asthma and the effects of prenatal air pollution exposure than females(12, 68, 69). This may be partly due to greater inflammatory signalling male fetal development(70, 71) and relatively slower large airway development in males compared to females(72). DNAm differences induced by air pollution that potentiate either of these effects could contribute to increase respiratory disease risk in males. In this study, prenatal PM_2.5_ exposure was associated with higher DNAm in the body of *SERPINB9* in males and lower DNAm in females. *SERPINB9* is expressed in several immune cell types(52) and inhibits intracellular granzyme B, protecting releasing cells from apoptosis(73). It also inhibits caspase-1, preventing maturation of IL-1β and inflammation(74, 75). The *SERPINB9* DMR identified here overlaps with a sex-specific DMR previously associated with forced expiratory volume in one second (FEV₁) in males(76). No associations were observed between PM_2.5_-induced *SERPINB9* DNAm differences and wheeze phenotypes in either males or females in this study. However, it remains plausible these DNAm differences could still contribute to sex-specific differences in lung function. Further replication is needed to clarify the role of *SERPINB9* DNAm in human respiratory health.

Most epigenome-wide studies of prenatal air pollution exposure have been conducted in primarily European birth cohorts(18–20, 77, 78). While these studies have provided meaningful insights into infant DNAm differences, their findings may have limited generalizability to more racially diverse populations. For example, we were unable to replicate many of the air pollution-associated DNAm findings from these European cohorts in the subset of CANDLE participants, in which over half (53%) of mothers identified as Black or African American. In contrast, several DNAm associations observed in PRISM(19), where a larger proportion of mothers identified as Hispanic (42%) or African American (18%), were replicated here in CANDLE. Shared regional, sociodemographic, or cultural differences between PRISM and CANDLE may underlie the greater consistency in findings, compared to those from predominantly European cohorts. These discrepancies underscore the need for greater diversity in DNAm and air pollution research and replication across sociodemographic populations should be conducted to ensure results are robust.

This study had several strengths and limitations. One advantage was the availability of prenatal air pollution estimates, which allowed us to compare DNAm alterations across multiple pollutant exposures(79). Another advantage of this study is the use of cord blood to investigate the role of DNAm changes in early-life wheeze phenotypes. While cord blood is commonly used due to its accessibility and relevance to prenatal exposures, it also enables investigation of DNAm in white blood cells, including T-cells, which contribute to immune dysfunction underlying wheeze and asthma(80). Genes highlighted in this study, including *HLA-DPA1/DPB1* and *SERPINB9*, are expressed in both cord blood and lung tissue(81), supporting the plausibility that observed DNAm differences could influence immune pathways relevant to respiratory disease. Further research is needed to replicate and characterize the role of DNAm differences in observed in this study on immune function and respiratory health and to determine if any of these differences are shared in lung tissue(82, 83). Future studies could consider animal models to interrogate the effects of different air pollution components on infant DNAm, and lung DNAm and function directly. For instance, a mouse model of prenatal cigarette exposures used by our lab successfully reproduced DNAm changes observed in human cord blood and also identified novel DNAm changes in lung tissue(71)

In terms of limitations, we used genetic ancestry PCs to describe population variation along a continuum and incorporated them in the statistical models to account for genetic variation within ancestral populations. While genetic ancestry PCs are often correlated with racial and ethnic identity, they are distinct from those social constructs and thus, cannot be considered as proxies of self-identified racial and ethnic identities (84). We did not include self-identified racial or ethnic identity in our models as we were interested in identifying DNAm alterations that were robust across populations. Therefore, we acknowledge that some of our associations may be influenced by residual confounding due to unaccounted racial and ethnic differences, or to ongoing racism and racist socioeconomic policies. Future studies thoughtfully designed to focus on these levels of inquiry may be of value(85). Finally, while this study includes more racial diversity than previous studies, CANDLE includes primarily Black/African American and White families. Replication of these findings across more populations is necessary before they can be generalized.

Together, the results from this study expand our understanding of how prenatal exposure to NO_2_, PM_2.5_, and PM_10_ affect DNAm. By leveraging the racially and socioeconomically diverse CANDLE cohort, we identified unique DNAm signatures associated with NO_2_, PM_2.5_, and PM_10_ exposure, and were able to replicate previous findings in cohorts with similar socioeconomic and racial structure. Importantly, the findings from this study highlight the potential for maternal diet micronutrients and overall diet quality to at least partially buffer some of these DNAm differences, supporting the role of a nutrient dense diet during pregnancy for child health.

## Data availability

Supplementary files are available for download on Zenodo (10.5281/zenodo.19240961). Code used for this analysis is available on Github at https://github.com/mjoneslab/CANDLE_cord_blood.

## Acknowledgments

The authors would like to acknowledge the participants of the CANDLE study.

## Notes

### Competing Interest Statement

The authors have declared no competing interest.

### Summary of Updates

The manuscript has been revised to include a Zenodo DOI link to download supplementary files and a link to the github repository housing analysis code.

### Funding

- Canadian Institutes of Health Research

