## Supporting Information (SI) for "Prenatal diet buffers infant epigenetic changes linked to pollution and transient wheeze"

### SI Methods

#### CANDLE study information

The Conditions Affecting Neurocognitive Development and Learning in Early Childhood (CANDLE) study was funded by the Urban Child Institute (UCI) to examine the effects of prenatal and early-life exposures and experiences on child health and development(1). University of Tennessee Health Science Centre (UTHSC) staff and clinicians conducted the study, including recruiting participants and collecting data. The study location of Shelby County, Tennessee was selected as it represents a racially diverse population with higher rates of adverse birth and health outcomes compared to the United States average. Mother-child dyads were considered eligible based on the following criteria: was a Shelby country resident, was between 16 and 40 years old, was between 16 and 28 weeks gestation, able to speak and understand English, planned to deliver at a participating healthcare center, had a singleton pregnancy, and considered to be low-risk (as defined by UCI) for pregnancy complications(1).

#### Wheeze phenotype definitions

Participants with transient wheeze were defined as an affirmative response to the question “Has your child ever had wheezing or whistling in the chest at any time in the past?” at either the year three or year four-six visit, and a dissenting response to the question “Has your child ever had wheezing or whistling in the chest in the last 12 months?” administered at the year four-six visit. Persistent wheeze was defined as an affirmative response to the question “Has your child ever had wheezing or whistling in the chest at any time in the past?” at both the year three and year four-six visit and an affirmative response to the question “Has your child ever had wheezing or whistling in the chest in the last 12 months?” administered at the year four-six visit. Late-onset wheeze was defined as a dissenting response to the question “Has your child ever had wheezing or whistling in the chest at any time in the past?” at the year three visit, and affirmative responses to the questions “Has your child ever had wheezing or whistling in the chest at any time in the past?” and “Has your child ever had wheezing or whistling in the chest in the last 12 months?” at the year four-six visit.

#### Genotyping and genetic ancestry calculations

Genotyping of CANDLE participant cord blood samples was performed using the Illumina Infinium Global Screening Array, followed by quality control in GenomeStudio where loci with GenCall scores <0.15, samples with call rates <0.97, samples and probes with GenCall rates <0.10, and probes with abnormal clustering were removed. Further filtering removed probes with a minor allele frequency <0.01, with heterozygote excess <-0.3 or >0.2, with normalized Hardy-Weinberg p-value $<10^{-16}$, or with short-range linkage disequilibrium. CANDLE participating genotyping data was combined with data for the five superpopulations in the 1000 Genomes projects and principal component analysis conducted to calculate genetic ancestry PCs. The first two genetic ancestry PCs accounting for >0.90 genetic variation and were included in linear regressions (**File S105**).

### Estimation of cord blood cell proportions

Cord blood is composed of several cell types, with each cell-type exhibiting a unique cell-specific DNA methylation (DNAm) signature. It is necessary to correct for differences in cell type proportions between samples to avoid spurious associations. Cord blood cell type proportions were not directly measured for CANDLE participants. Therefore, we applied a reference-based method to estimate cord blood mononuclear cell (CBMC) proportions in each sample. Specifically, we used the Illumina MethylationEPIC array cord blood reference set from the *FlowSorted.CordBloodCombined.450k* R package and the *estimateCellCounts()* function from the *minfi* R package to estimate the relative proportions of 7 cell types (B-cells, CD4 T-cells, granulocytes, monocytes, NK cells, and nucleated red blood cells) in CBMC samples. A small offset of 0.0001 was added to cell type estimates to remove the implausible scenario where a cell type was estimated as zero.

Raw estimations of cell type proportions should not be directly included in linear regressions as their compositional nature may result in multicollinearity. Principal component analysis (PCA) can remove these compositional constraints; however, a transformation must first be performed to make compositional cell type estimates suitable for PCA. We performed isometric-log transformation on CBMC estimates followed by robust PCA using the *robCompositions* R package. The first four principal components (PCs) accounting for >90% variation in CMBC estimations were included in robust multivariable linear models (**File S106**).

### Model covariates

Several *a priori* selected covariates were included in linear regressions to account for biological variation including participant biological sex and gestational age. Maternal characteristics including prenatal smoking and household income were assessed by CANDLE staff via a questionnaire administered at the baseline visit (16 to 26 weeks gestation)(1). Maternal prenatal tobacco use was considered as an affirmative response to the question “Are you currently using tobacco?”. CANDLE staff asked participants to estimate their total annual household on a categorical scale. Estimated annual household income was used to calculate adjusted household income, which accounts for the number of children (<18 years old) and adults (>=18 years old) in the home. Participant information including biological sex and gestational age was obtained from medical records by registered nurse with training in obstetrics(1). Each model also included four surrogate variables generated separately for $NO_2$, $PM_{2.5}$, and $PM_{10}$ by the SVA package to remove residual technical variation effects(2)(**Files S107-S109**).

**SI Figures**

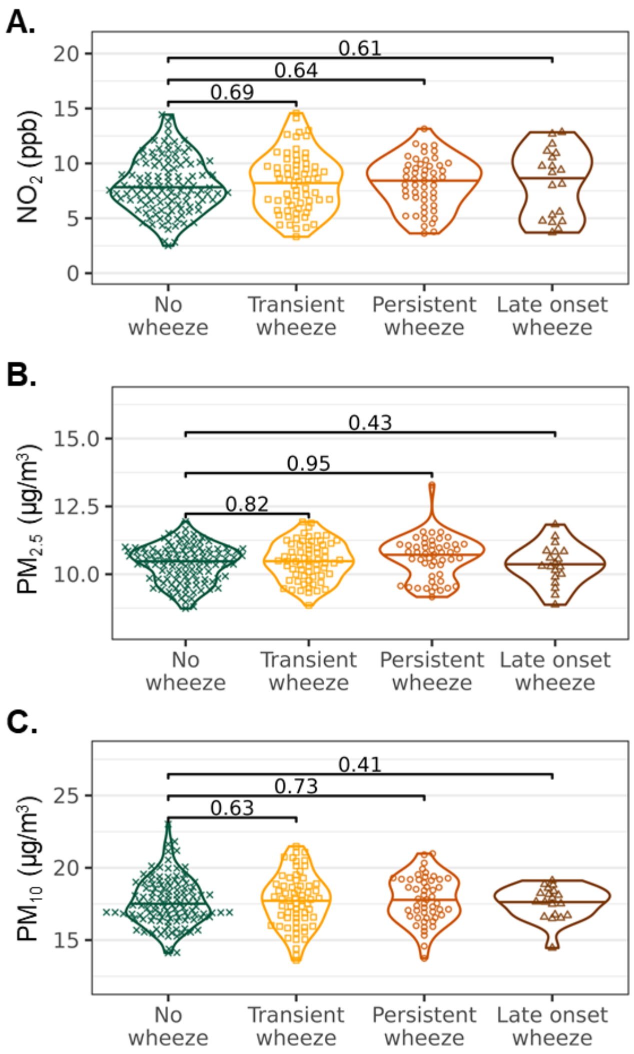


**Fig. S1. Relationship between prenatal $NO_2$, $PM_{2.5}$, and $PM_{10}$ exposures and never (N=119), transient (N=65), persistent (N=51), or late onset (N=19) wheeze.** Prenatal (**A**) $NO_2$ exposures, (**B**) $PM_{2.5}$ exposures, and (**C**) $PM_{10}$ exposures were not significantly different between individuals without wheeze and in those with transient, persistent, or late onset wheeze. Mean and standard deviations of $NO_2$ exposure were 8.0±2.5 ppb, 8.3±2.7 ppb, 8.2±2.3 ppb, and 8.3±3.1 ppb; mean and standard deviations of $PM_{2.5}$ exposures were 10.4±0.7 µg/m$^3$, 10.5±0.7 µg/m$^3$, 10.6±0.7 µg/m$^3$, and 10.4±0.7 µg/m$^3$; and mean and standard deviations of $PM_{10}$ exposure were. 17.7±1.6 µg/m$^3$, 17.7±1.8 µg/m$^3$, 17.8 ±1.6 µg/m$^3$, 17.6±1.1 µg/m$^3$ in individual without wheeze or with transient, persistent, or late-onset wheeze, respectively. One sided t-tests investigated whether prenatal air pollutant exposures were higher in individuals with wheeze compared to individuals without wheeze.

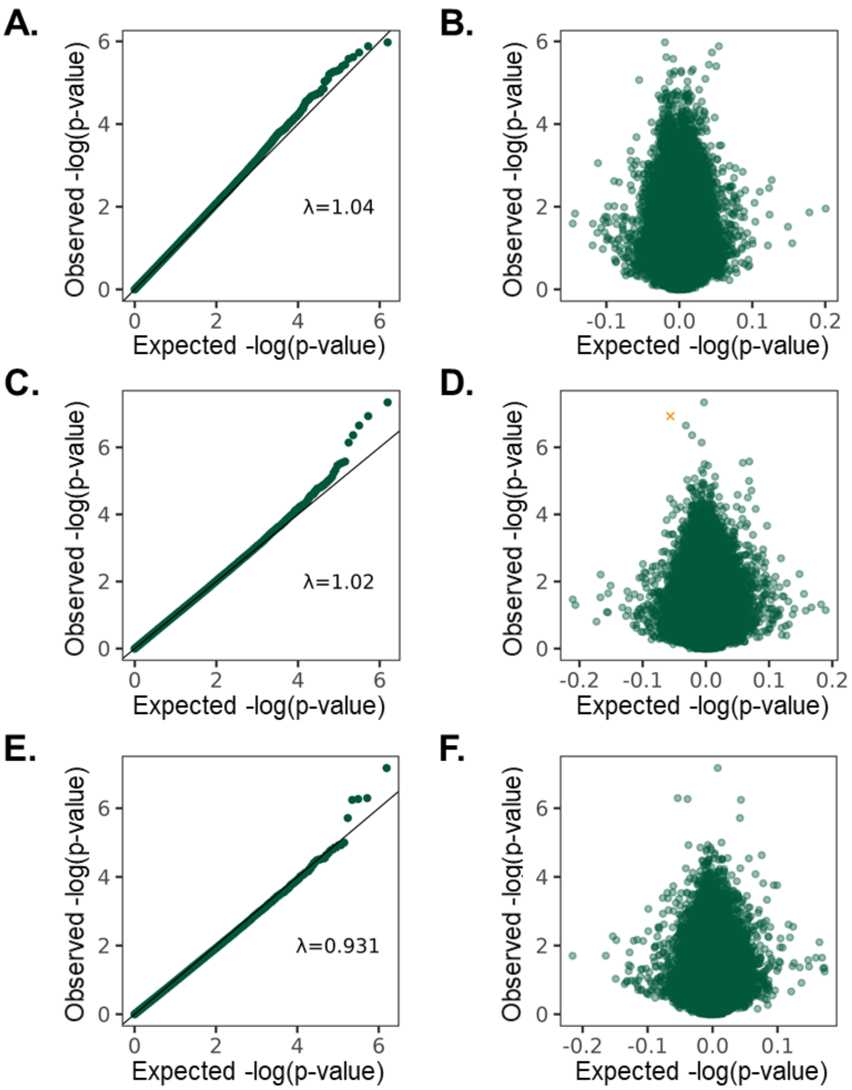


**Fig. S2. Model fits of the epigenome-wide analyses of prenatal $NO_2$, $PM_{2.5}$, and $PM_{10}$ in cord blood DNA methylation (DNAm) of CANDLE participants (N=515).** (**A**) The model of prenatal $NO_2$ exposure exhibited a genomic inflation of 1.04. (**B**) No individual CpGs exhibited statistically significant alterations in DNAm effect size associated with prenatal $NO_2$ exposure at 5% FDR. (**C**) The model of prenatal $PM_{2.5}$ exposure exhibited a genomic inflation of 1.02. (**D**) DNAm at one CpG was significantly and negatively (adjusted p=0.046, effect size=-0.06) associated with prenatal $PM_{2.5}$ exposure. (**E**) The model of prenatal $PM_{10}$ exposure exhibited a genomic inflation of 0.93. (**F**) No individual CpGs exhibited statistically significant alterations in DNAm effect size associated with prenatal $PM_{10}$ exposure at 5% FDR.

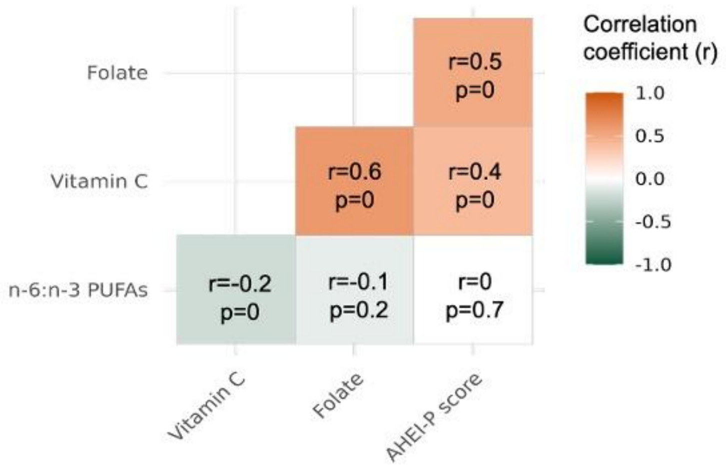


**Fig. S3. Correlations between maternal intake of vitamin C, polyunsaturated fatty acids PUFAs), and folate and the Alternative Health Eating Index Pregnancy (AHEI-P) score (N=444).** Significant ($p<0.05$) Pearson correlations (r) were identified between several micronutrients and AHEI-P score.

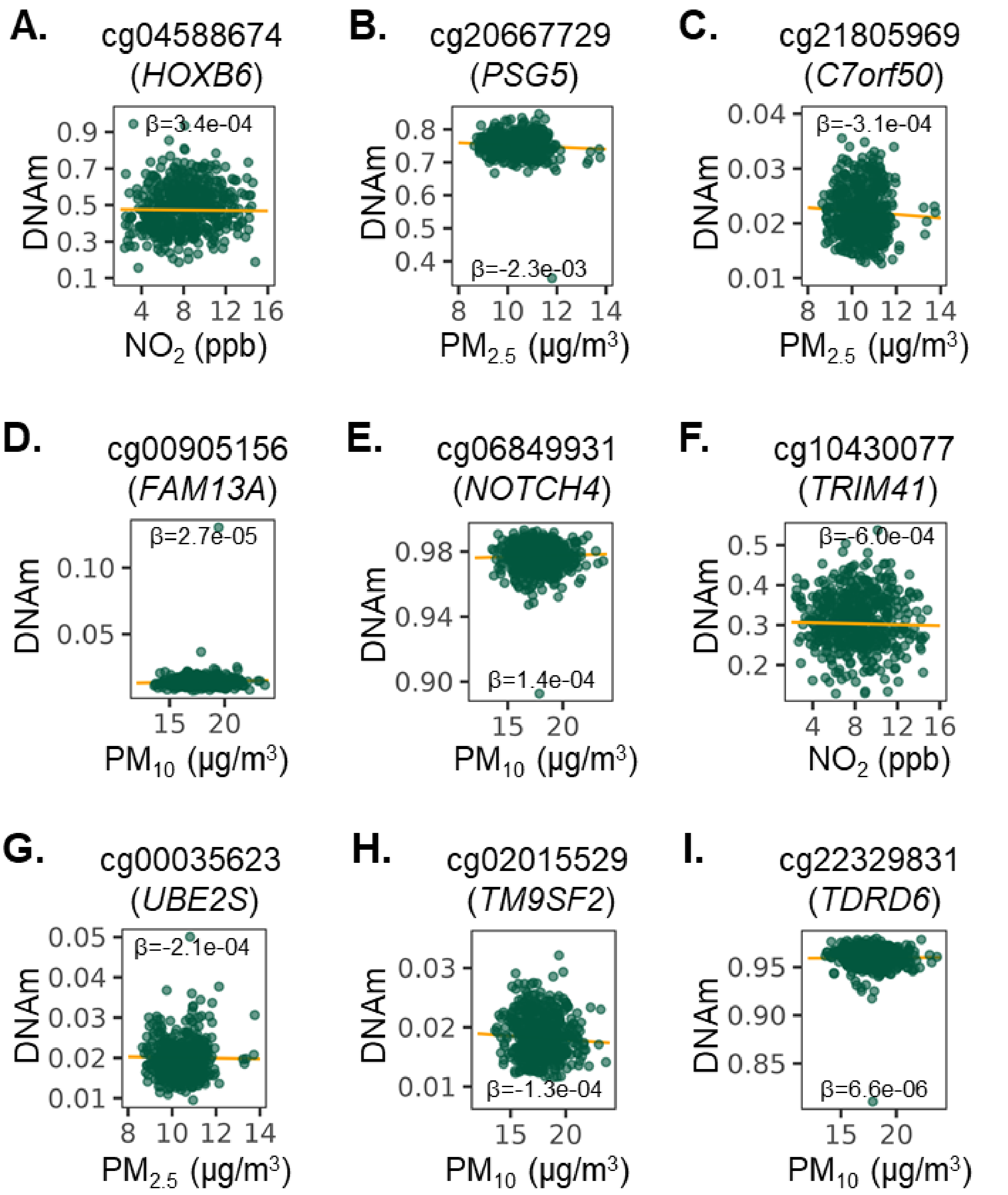


**Fig. S4 Replication of DNA methylation (DNAm) changes identified in previous epigenome-wide studies of cord blood and prenatal $NO_2$, $PM_{2.5}$, or $PM_{10}$ exposure in CANDLE participants (N=515).** DNAm changes associated with prenatal air pollution exposures at (**A**) cg04588674, (**B**) cg20667729, (**C**) cg21805969, (**D**) cg00905156, and (**E**) cg06849931 replicated in CANDLE participants, while DNAm changes at (**F**) cg10430077, (**G**) cg00035623, (**H**) cg02015529, and (**I**) cg22329831 did not. β represents the coefficient of prenatal air pollutant exposures obtained from robust linear regression of DNAm β-values.

**SI Tables**

**Table S1.** Study population (N=515) characteristics stratified by biological sex.

| | Males (N=249) | Females (N=266) |
|---|---|---|
| **Prenatal $NO_2$ (ppb) [mean±sd]** | 8.2 ± 2.5 | 8.1 ± 2.7 |
| **Prenatal $PM_{2.5}$ (μg/$m^3$) [mean±sd]** | 10.5 ± 0.8 | 10.4 ± 0.8 |
| **Prenatal $PM_{10}$ (μg/$m^3$) [mean±sd]** | 17.7 ± 1.7 | 17.8 ± 1.7 |
| **Maternal self-identified race** | | |
| Black/African American | 134 (54%) | 140 (53%) |
| White | 99 (40%) | 106 (40%) |
| Asian | 3 (1%) | 3 (1%) |
| American Indian/Alaska Native | 1 (0%) | 0 (0%) |
| Other | 2 (1%) | 1 (0%) |
| Multiple Race | 10 (4%) | 16 (6%) |
| **Gestational Age (weeks) [mean±sd]** | 39 ± 1 | 39 ± 1 |
| **Adjusted income (USD) [mean±sd]** | 23400 ± 18100 | 21800 ± 17600 |
| **Maternal prenatal tobacco use [count(%)]** | | |
| Yes | 13 (5%) | 28 (11%) |
| No | 236 (95%) | 238 (89%) |
| **Transient wheeze at age four to six [count(%)]** | | |
| Yes | 34 (14%) | 31 (12%) |
| No | 166 (67%) | 189 (71%) |
| Missing | 49 (20%) | 46 (17%) |
| **Persistent wheeze at age four to six [count(%)]** | | |
| Yes | 29 (12%) | 22 (8%) |
| No | 175 (70%) | 216 (81%) |
| Missing | 45 (18%) | 28 (11%) |
| **Late onset wheeze at age four to six [count(%)]** | | |
| Yes | 9 (4%) | 10 (4%) |
| No | 182 (73%) | 209 (79%) |
| Missing | 58 (23%) | 47 (18%) |
| **Ever atopy at age four to six [count(%)]** | | |
| Yes | 64 (26%) | 65 (24%) |
| No | 131 (53%) | 152 (57%) |
| Missing | 54 (22%) | 49 (18%) |
| **Current asthma at age four to six [count(%)]** | | |
| Yes | 38 (15%) | 27 (10%) |
| No | 129 (52%) | 149 (56%) |
| Missing | 82 (33%) | 90 (34%) |
| **Vitamin C dietary density (mg/kcal) [mean±sd]** | 76.8 ± 30.1 | 72.2 ± 31.1 |

| | | |
|---|---|---|
| Missing [count(%)] | 10 (4.0%) | 17 (6.4%) |
| **PUFA dietary density (gms/kcal) n6:n3 ratio [mean±sd]** | 9.1 ± 1.3 | 9.0 ± 1.3 |
| Missing [count(%)] | 10 (4.0%) | 17 (6.4%) |
| **Folate dietary density (mcg/kcal) [mean±sd]** | 147 ± 57.3 | 139 ± 50.8 |
| Missing [count(%)] | 10 (4.0%) | 17 (6.4%) |
| **Alternative Healthy Eating Index Pregnancy (AHEI-P) [mean±sd]** | 62.7 ± 12.0 | 59.2 ± 12.6 |
| Missing [count(%)] | 10 (4.0%) | 17 (6.4%) |
| **Estimated cord blood cell proportions [mean±sd]** | | |
| B-Cells | 0.08 ± 0.04 | 0.07 ± 0.04 |
| CD4 T-cells | 0.15 ± 0.06 | 0.14 ± 0.06 |
| CD8 T-cells | 0.11 ± 0.04 | 0.09 ± 0.03 |
| Granulocytes | 0.43 ± 0.13 | 0.45 ± 0.13 |
| Monocytes | 0.12 ± 0.04 | 0.12 ± 0.04 |
| NK cells | 0.04 ± 0.03 | 0.04 ± 0.03 |
| nRBCs | 0.12 ± 0.07 | 0.12 ± 0.10 |

**Table S2.** Study population characteristics stratified by health outcome.

| | Unaffected healthy control (N=119) | Transient wheeze (N=65) | Persistent wheeze (N=51) | Late onset wheeze (N=19) |
|---|---|---|---|---|
| **Prenatal $NO_2$ (ppb) [mean±sd]** | 8.1 ± 2.5 | 8.3 ± 2.7 | 8.2 ± 2.3 | 8.3 ± 3.1 |
| **Prenatal $PM_{2.5}$ (μg/$m^3$) [mean±sd]** | 10.4 ± 0.7 | 10.5 ± 0.7 | 10.6 ± 0.8 | 10.4 ± 0.7 |
| **Prenatal $PM_{10}$ (μg/$m^3$) [mean±sd]** | 17.7 ± 1.6 | 17.7 ± 1.8 | 17.8 ± 1.6 | 17.6 ± 1.1 |
| **Maternal self-identified race [count(%)]** | | | | |
| Black/African American | 54 (45%) | 33 (51%) | 37 (73%) | 7 (37%) |
| White | 54 (45%) | 28 (43%) | 10 (20%) | 10 (53%) |
| Asian | 1 (1%) | 0 (0%) | 0 (0%) | 0 (0%) |
| American Indian/Alaska Native | 0 (0%) | 0 (0%) | 0 (0%) | 0 (0%) |
| Other | 1 (1%) | 0 (0%) | 0 (0%) | 1 (5%) |
| Multiple Race | 9 (8%) | 4 (6%) | 4 (8%) | 1 (5%) |
| **Gestational Age (weeks) [mean±sd]** | 39 ± 1 | 39 ± 1 | 39 ± 2 | 40 ± 1 |
| **Adjusted income (USD) [mean±sd]** | 26600 ± 18700 | 23500 ± 17200 | 16700 ± 14700 | 23100 ± 17800 |
| **Maternal prenatal tobacco use [count(%)]** | | | | |
| Yes | 10 (8%) | 3 (5%) | 2 (4%) | 2 (11%) |
| No | 109 (92%) | 62 (95%) | 49 (96%) | 17 (89%) |
| **Transient wheeze at age four to six[count(%)]** | | | | |
| Yes | 0 (0%) | 65 (100%) | 0 (0%) | 0 (0%) |
| No | 119 (100%) | 0 (0%) | 51 (100%) | 19 (100%) |
| **Persistent wheeze at age four to six [count(%)]** | | | | |
| Yes | 0 (0%) | 0 (0%) | 51 (100%) | 0 (0%) |
| No | 119 (100%) | 65 (100%) | 0 (0%) | 19 (100%) |
| **Late onset wheeze at age four to six [count(%)]** | | | | |
| Yes | 0 (0%) | 0 (0%) | 0 (0%) | 19 (100%) |
| No | 119 (100%) | 65 (100%) | 51 (100%) | 0 (0%) |
| **Ever atopy at age four to six [count(%)]** | | | | |
| Yes | 0 (0%) | 28 (43%) | 23 (45%) | 7 (37%) |

| | | | | |
|---|---|---|---|---|
| No | 119 (100%) | 37 (57%) | 26 (51%) | 12 (63%) |
| Missing | 0 (0%) | 0 (0%) | 2 (4%) | 0 (0%) |
| **Current asthma at age four to six [count(%)]** | | | | |
| Yes | 0 (0%) | 4 (6%) | 42 (82%) | 11 (58%) |
| No | 119 (100%) | 61 (94%) | 7 (14%) | 7 (37%) |
| Missing | 0 (0%) | 0 (0%) | 2 (4%) | 1 (5%) |
| **Vitamin C dietary density (mg/kcal) [mean±sd]** | 73.2 (28.8) | 76.2 (29.1) | 79.0 (33.9) | 70.3 (24.4) |
| Missing [count(%)] | 7 (5.9%) | 2 (3.1%) | 2 (3.9%) | 0 (0%) |
| **PUFA dietary density (gms/kcal) n6:n3 ratio [mean±sd]** | 9.2 ± 1.2 | 9.2 ± 1.4 | 9.0 ± 1.1 | 9.1 ± 1.4 |
| Missing [count(%)] | 7 (5.9%) | 2 (3.1%) | 2 (3.9%) | 0 (0%) |
| **Folate dietary density (mcg/kcal) [mean±sd]** | 148 ± 51.2 | 153 ± 54.7 | 131 ± 53.5 | 141 ± 42.2 |
| Missing [count(%)] | 7 (5.9%) | 2 (3.1%) | 2 (3.9%) | 0 (0%) |
| **Alternative Healthy Eating Index Pregnancy (AHEI-P) [mean±sd]** | 61.1 ± 11.1 | 65.5 ± 11.3 | 61.1 ± 13.8 | 64.0 ± 12.2 |
| Missing [count(%)] | 7 (5.9%) | 2 (3.1%) | 2 (3.9%) | 0 (0%) |
| **Estimated cord blood cell proportions [mean±sd]** | | | | |
| B-Cells | 0.07 ± 0.03 | 0.08 ± 0.04 | 0.08 0.05 | 0.07 ± 0.03 |
| CD4 T-cells | 0.14 ± 0.06 | 0.15 ± 0.07 | 0.13 ± 0.05 | 0.15 ± 0.05 |
| CD8 T-cells | 0.10 ± 0.03 | 0.10 ± 0.04 | 0.10 ± 0.03 | 0.10 ± 0.05 |
| Granulocytes | 0.46 ± 0.12 | 0.43 ± 0.14 | 0.43 ± 0.13 | 0.43 ± 0.13 |
| Monocytes | 0.12 ± 0.03 | 0.12 ± 0.04 | 0.12 ± 0.05 | 0.10 ± 0.04 |
| NK cells | 0.04 ± 0.03 | 0.04 ± 0.03 | 0.04 ± 0.03 | 0.04 ± 0.02 |
| nRBCs | 0.12 ± 0.09 | 0.13 ± 0.09 | 0.15 ± 0.10 | 0.16 ± 0.14 |
